# Mechanism of 30S subunit recognition and modification by the conserved bacterial ribosomal RNA methyltransferase RsmI

**DOI:** 10.1101/2025.08.28.672957

**Authors:** Mohamed I. Barmada, Erin N. McGinity, Suparno Nandi, Debayan Dey, Natalia Zelinskaya, George M. Harris, Lindsay R. Comstock, Christine M. Dunham, Graeme L. Conn

## Abstract

Ribosomal RNA (rRNA) modifications are important for ribosome function and can influence bacterial susceptibility to ribosome-targeting antibiotics. The universally conserved 16S rRNA nucleotide C1402, for example, is the only 2’-*O*-methylated nucleotide in the bacterial small (30S) ribosomal subunit and this modification fine tunes the shape and structure of the peptidyl tRNA binding site. The Cm1402 modification is incorporated by the conserved bacterial 16S rRNA methyltransferase RsmI, but it is unclear how RsmI is able to recognize its 30S substrate and specifically modify its buried target nucleotide. We determined a 2.42 Å resolution cryo-EM structure of the RsmI-30S complex and, with accompanying functional analyses, show that RsmI anchors itself to the 30S subunit through multiple contacts with a conserved 16S rRNA surface previously only seen in the assembled subunit. This positions RsmI to induce an extensive h44 distortion to access C1402 that is unprecedented among 16S rRNA methyltransferases characterized to date. These analyses also reveal an essential contribution to 30S subunit interaction made by the previously structurally uncharacterized RsmI C-terminal domain, RsmI-induced RNA-RNA interactions with C1402, and an unappreciated dependence on a divalent metal ion for activity that suggests RsmI may be first of a distinct class of metal- and SAM-dependent RNA *O*-methyltransferases. This study significantly expands our mechanistic understanding of how intrinsic bacterial methyltransferases like RsmI modify their rRNA targets. Further, recognition of distant ribosome features and extensive unfolding of a critical rRNA functional center point to a potential role in accurate 30S subunit biogenesis.

## Introduction

One hallmark of accurate bacterial ribosome biogenesis is methylation of select nucleotides at the functional centers of both ribosomal subunits (1). These modified nucleotides play important roles in the process of individual subunit assembly (2), translational fidelity (3, 4), ribosome-ligand interactions (5), and antibiotic susceptibility (6). Among the conserved methylated nucleotides in the small (30S) ribosomal subunit, C1402 is the only 16S rRNA nucleotide to be methylated at both its nitrogenous base (on the N4 atom to form m^4^C1402) and its ribose 2’-OH (to form Cm1402). The conserved S-adenosyl-L-methionine (SAM)-dependent methyltransferase RsmI is responsible for incorporating the latter modification (7), which is the only conserved ribose 2’-O-methyl modification in the bacterial 30S subunit.

The dual methylated C1402 (m^4^Cm1402) is located within helix 44 (h44), forming a non-canonical base pair with A1500 (8). m^4^Cm1402 plays important roles in fine tuning the shape and structure of the 30S subunit peptidyl tRNA binding site (P site) and makes a direct contact with the mRNA codon (4, 9). Loss of RsmI, and thus the Cm1402 modification, results in defects in translational fidelity, including an increase in stop codon readthrough, a decrease in mRNA reading frame maintenance (7), and increased sensitivity to certain antibiotics (6). The level of C1402 2’-O-methylation appears dynamically regulated in response to antibiotics (10) and also contributes to sensitivity to other stresses such as oxidative stress (11). While much is known about the functional impacts of the Cm1402 modification, how the ribose methylation is incorporated by RsmI is currently less well understood.

RsmI functions as a dimer to incorporate the Cm1402 modification within the assembled 30S subunit but is unable to modify naked 16S rRNA or 70S ribosomes *in vitro* (7, 12). These observations imply that RsmI likely acts on the 30S subunit late in biogenesis but before 70S ribosome assembly, where h44 becomes further occluded by the 50S subunit. While RsmI therefore presumably relies on conserved features of the assembled 30S subunit or late assembly intermediate for substrate recognition, the enzyme must also be able to manipulate the folded architecture of the 16S rRNA h44 to access the relatively deeply buried C1402 residue. However, why RsmI acts as a dimer and requires the mature or near mature 30S subunit as its substrate, and how RsmI distorts the h44 structure to access C1402 for modification are currently unknown.

To uncover the molecular basis behind C1402 recognition and modification by RsmI, we determined the structure of the *Escherichia coli* RsmI-30S subunit complex to a global resolution of 2.42Å using cryogenic electron microscopy (cryo-EM) complemented by functional studies of key RsmI residues. These studies reveal that RsmI anchors itself to the 30S subunit by binding to a highly conserved surface of 16S rRNA helices previously only seen in the assembled 30S subunit, relying on both protomers of its dimeric structure for distinct but equally essential roles in substrate recognition. One RsmI protomer extensively distorts h44 to access C1402, while the other recognizes a distant structural feature of the 30S subunit via a functionally critical but previously structurally undefined C-terminal domain (CTD). These studies additionally reveal catalysis of C1402 methylation by RsmI to be a divalent metal ion-dependent process, a unique feature among 16S rRNA methyltransferases characterized to date. Overall, this work reveals an extraordinarily high degree of rRNA manipulation by an rRNA methyltransferase and provides key insights into both the mechanism of action of intrinsic ribosomal 16S rRNA methyltransferases as well as pointing to their potential contributions to 30S subunit biogenesis.

## Results

### Structure of the RsmI-30S subunit complex

The 30S subunit lacking the Cm1402 modification (“30S-Δ*rsmI*”) was isolated from an *E. coli* Δ*rsmI* strain and confirmed as a suitable substrate for RsmI using an *in vitro* methyltransferase assay (**Supplementary Fig. S1**). The RsmI-30S complex was prepared by mixing 30S-Δ*rsmI*, purified recombinant RsmI and the SAM analog 5’-(diaminobutyric acid)-N-iodoethyl-5’-deoxyadenosine ammonium hydrochloride (“NM6”) in a 1:3:30 ratio. Through the action of RsmI, NM6 becomes covalently attached to the substrate in an alkylation reaction (13, 14), trapping the enzyme-30S substrate complex in an immediately post-catalytic state by virtue of the enzyme’s affinity for both substrate and co-substrate analog.

With this complex, we used single-particle cryo-EM to generate a map of the entire RsmI-30S subunit complex as well as a focused map of RsmI and its adjacent 16S rRNA binding interface on the 30S subunit to global resolutions of 2.42 Å and 2.55 Å, respectively (**Fig. 1A,B**; **Supplementary Figs. S2**, **S3A,B**, and **Table S1**). Together, these maps enabled the full modeling of the structure of the RsmI-30S complex (**Fig. 1C**). Notably, the higher local resolution of the focused map (**Supplementary Fig. S3C,D**) allowed for confident modeling of the C1402 ribose 2’-O-linked NM6 within the RsmI SAM binding site (**Supplementary Fig. S3E**), the extensive interactions between RsmI and the 16S rRNA (**Supplementary Fig. S3F-H**), and the most complete structure of the RsmI dimer, to date, including the previously structurally uncharacterized CTD (**Fig. 1D**).

**Figure 1.**
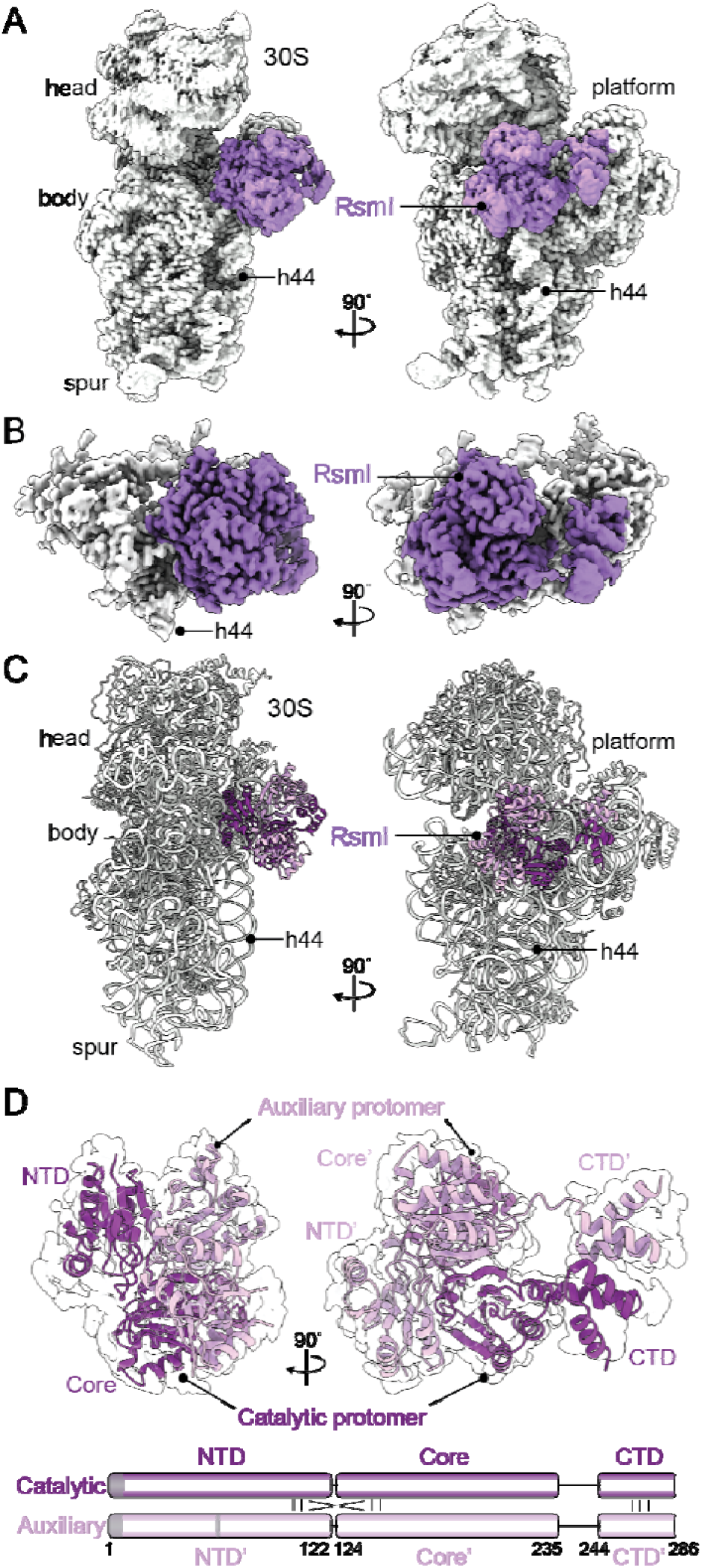
Structure of the RsmI-30S subunit complex. (*A*) Sharpened cryo-EM map of the RsmI-30S complex. (*B*) Sharpened cryo-EM map of RsmI and its 16S rRNA binding surface on the 30S sub nit. (*C*) Model of the RsmI-30S complex. (*D*) Model of the RsmI dimer including the newly resolved CTD overlaid on the map and accompanied by cartoon representation of the RsmI domain organization and interaction sites between the two protomers (below). Regions absent in the final RsmI model are shaded in gray. Domains (and α helices) of the auxiliary RsmI protomer are denoted with a prime (‘) in this and subsequent figures.

Overall, the structure reveals that RsmI binds to an rRNA surface comprising h44 and the 30S platform (**Fig.1A,C**), employing its three major domains for distinct functions: an N-terminal domain (NTD) which makes extensive contacts with 16S rRNA, a core domain which also contacts 16S rRNA and contains a Rossmann-like fold with SAM binding and methyltransferase catalytic activities, and the newly modeled CTD connected to the core domain by a short, flexible linker that mediates contacts most distant from the target nucleotide (**Fig. 1D**). Interaction of RsmI with the 30S subunit additionally distinguishes the two protomers within the dimeric structure, with one carrying the NM6 that is covalently attached to C1402 and thus anchored on h44 (which we refer to as the “catalytic” protomer) and the second (“auxiliary” protomer) in the opposite orientation with its SAM binding pocket facing away from h44 toward the solvent (**Fig. 1C,D**).

### RsmI severely distorts h44 and positions C1402 for methylation using an array of conserved positive and aromatic residues

C1402 is located deep in the minor groove of h44 with its ribose 2’-OH group facing toward the interior of the 30S subunit body, making the RsmI target site inaccessible in the assembled subunit. As such, we anticipated that a major rearrangement of h44 RNA would be necessary to allow access for modification by RsmI. Indeed, our structure reveals that RsmI dramatically disrupts h44 surrounding C1402, displacing nucleotides on both strands and breaking a total of 7 base pair interactions, and additionally displacing nucleotides A1502-U1506 which connect h44 and h45 (**Fig. 2A**). The 5’-strand of h44 containing C1402 (nucleotides C1399-A1408) is pulled outward from its usual position to make multiple contacts with the NTD and Core domain of the catalytic protomer of RsmI. In contrast, the complementary nucleotides of the 3’-strand (nucleotides G1494-C1501), do not make extensive interactions with RsmI and appear highly dynamic. The backbone path of these nucleotides was, however, visible using a lower resolution map of the full complex, allowing for approximate modelling of their location (**Supplementary Fig. S4A**). To confirm that the disruption of h44 is due to binding of RsmI and not intrinsic to the hypomethylated 30S-Δ*rsmI*, we isolated a minor subset of particles from our dataset that had almost no observed RsmI occupancy, resulting in a 2.91 Å map from which we determined the structure of the free 30S-Δ*rsmI* (**Supplementary Fig. S4B**). Superimposing the structure of this free 30S-Δ*rsmI* to the wild-type 30S subunit revealed the structures to be almost identical, with no h44 deformations and C1402 occupying its canonical position.

**Figure 2.**
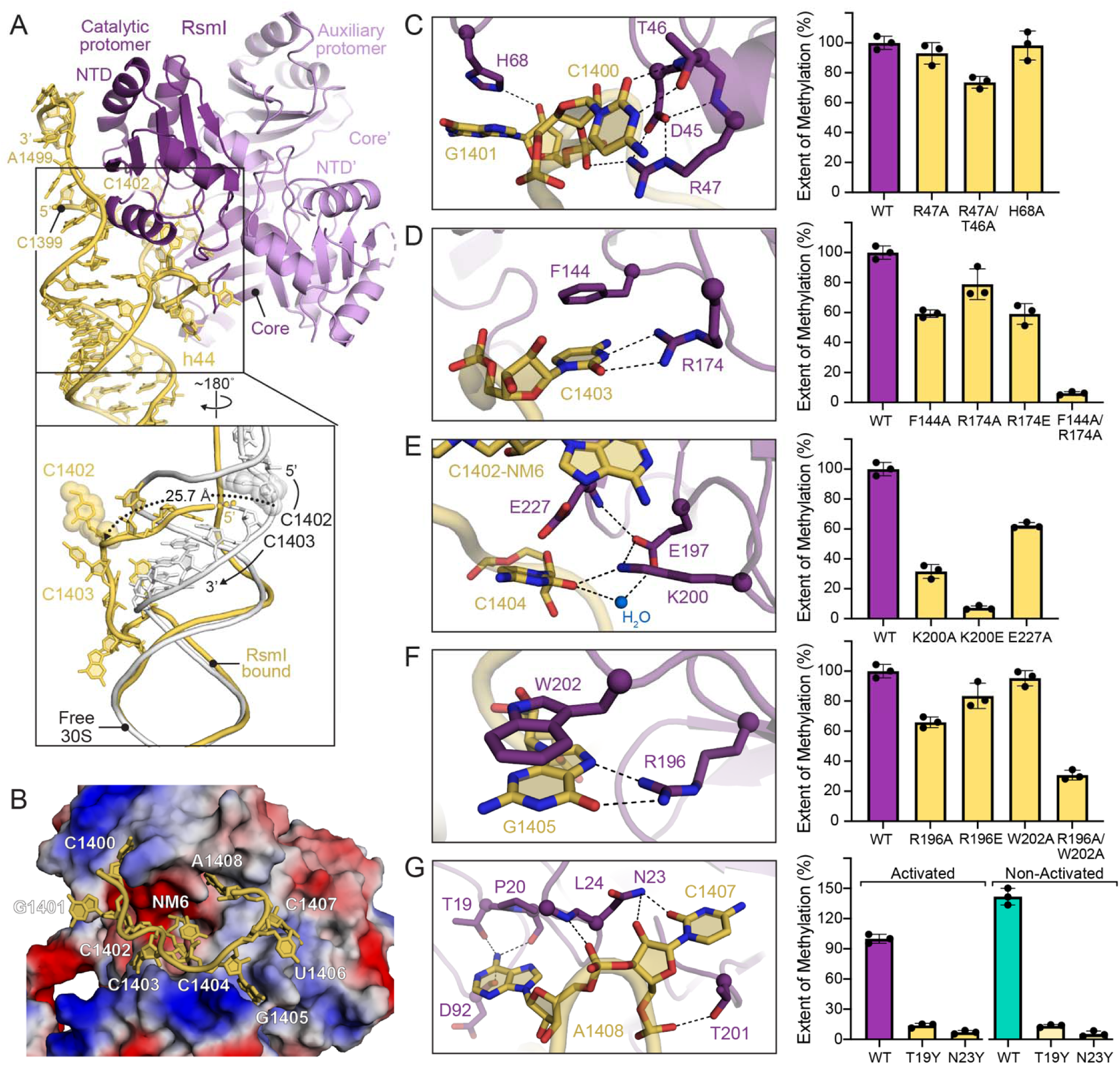
The RsmI catalytic protomer manipulates h44 to access target nucleotide C1402. (*A*) View of the RsmI-h44 interface with a comparison of h44 residues C1400-A1408 between RsmI-bound (gold) and free 30S subunit (white). In the zoomed view, the approximate path of the rRNA backbone of the h44 strand (5’ of C1402) is indicated with gold spheres as this region could not be modeled due to disorder. (*B*) RsmI electrostatic surface with h44 residues C1400-A1408. Zoomed views of the interaction networks between RsmI and h44 nucleotides (*left*) and activity analyses for RsmI variants (*right*) for: (*C*) C1400 and G1401, (*D*) C1403, (*E*) C1404, (*F*) G1405, and (*G*) C1407 and A1408.

To unwind h44 and access C1402, RsmI surrounds the nucleotides before (C1400 and G1401) and after C1402 (C1403 to A1408) with a collection of conserved aromatic and positive residues derived exclusively from the catalytic protomer (**Fig. 2B**; **Supplementary Fig. S5** and **Table S2**). These residues form several interaction networks with the 16S rRNA nucleotides surrounding C1402 to disrupt the base stacking and base pairing interactions in which these nucleotides normally engage (**Fig. 2C-G**).

On the 5’-side of C1402, RsmI uses NTD residues D45, T46, R47, and H68 to form an interaction network surrounding C1400 and G1401 (**Fig. 2C**). However, despite their high conservation (**Supplementary Fig. S5B** and **Table S2**), individual substitution of R47 or H68 does not significantly reduce RsmI activity *in vitro* (**Fig. 2C**). Further, a double T46A/R47A substitution results in only a modest (∼35%) reduction, suggesting that interactions with 16S rRNA nucleotides that are 5’ of C1402 are not essential for RsmI to access and modify its target nucleotide. In contrast, the RsmI residue interaction networks surrounding nucleotides C1403, C1404, and G1405 on the 3’-side of C1402 appear collectively crucial for the ability of RsmI to access its target nucleotide (**Fig. 2D-F**). At C1403 and G1405, RsmI employs a similar strategy, surrounding each nucleotide with an aromatic residue (F144 and W202, respectively) that stacks with the nucleotide and a basic residue (R174 and R196, respectively) that hydrogen bonds with the base Watson-Crick face (**Fig. 2D,F**). While single substitutions of these RsmI residues only modestly impacted activity, double substitutions (F144A/R174A and R196A/W202A) resulted in more substantial reductions in C1402 methylation (**Fig. 2D,F**), highlighting the importance of these networks. At C1404, RsmI residue K200 also appears crucial in 16S rRNA interaction and balancing the negative charges present in the interaction network surrounding it. Loss of the K200 positive side chain (K200A) or reversal of its charge (K200E) significantly lowers or eliminates RsmI activity (**Fig. 2E**). Further away from C1402 on the 3’-side, nucleotide C1407 is surrounded by RsmI residues N23 and T201, while nucleotide A1408 is sequestered in a pocket formed by RsmI residues T19, P20, and D92 (**Fig. 2G**). Replacement of N23 and T19 with bulky tyrosine residues to prevent C1407 and A1408, respectively, from moving to occupy their new positions completely blocked the ability of RsmI to methylate C1402 (**Fig. 2G**).

In the bacterial cell, the mature 30S can interconvert between an “active” conformation in which h44 is in its canonical position with respect to the 30S subunit body and a predominant “inactive” conformation in free 30S subunits in which h44 is moved outward and the decoding center is not fully formed (15–17). As the process of 30S subunit isolation produces a mixed population containing both 30S species, we used an established 30S subunit activation process to ensure consistent, homogenous species were used in the studies described to this point. However, in light of the ability of RsmI to extensively unwind h44, we asked whether omitting the activation step might result in a more preferred 30S subunit substrate. Indeed, comparison of RsmI activity on activated and non-activated 30S subunits revealed a greater extent of methylation with non-activated 30S substrate over the full reaction time course (**Supplementary Fig. S6A**). In further experiments using a range of 30S subunit substrate concentrations, at both early and late time points, this difference was maintained but reduced in the latter (**Supplementary Fig. S6B,C**). Critically, however, the RsmI variants with amino acid substitutions (T19Y and N23Y) that disrupt interaction with h44 at C1407 and A1408, most distant from C1402, are equally unable to modify the non-activated 30S subunit (**Fig. 2G**). The extensive unfolding of h44 to access C1402 for modification thus appears essential regardless of the activation state of the 30S subunit.

These data collectively indicate that the unwinding of h44 depends on the coordinated movement of each nucleotide from A1408 to C1403, as disruption of the RsmI interaction network around any one of these nucleotides significantly impedes C1402 methylation (**Fig. 2D-G**). In contrast, disruption of RsmI contact with the nucleotides 5’ of C1402 (i.e., C1400 and G1401) had only minimal impact on its activity (**Fig. 2C**). Together, these observations suggest a directionality to the unwinding of h44 by RsmI, beginning with a major repositioning of A1408, which is likely to be accessible due to its position opposite the dynamic nucleotides A1492/A1493, and proceeding in a 3’ to 5’ direction along the 5’-strand of h44 to C1402. In this way, RsmI accesses C1402 by using its catalytic protomer to unwind and severely distort h44, thereby partitioning residues adjacent to C1402 away from their usual interactions through a redundant set of interaction networks.

### RsmI engages a conserved 16S rRNA surface to recognize its 30S subunit substrate

RsmI requires an assembled 30S subunit as its minimal substrate for C1402 modification, with its lack of activity on free 16S rRNA (7) indicating that some characteristic feature(s) only present in the mature (or near mature) 30S subunit are essential. Our structure reveals that in addition to the extensive interactions with h44, RsmI engages three additional 16S rRNA helices on the 30S subunit platform: h23b, h24a, and h45. These four 16S rRNA helices are distant in the primary sequence but are brought together in the assembled 30S subunit to form highly a conserved rRNA surface for RsmI-30S subunit interaction (**Fig. 3A, B**).

**Figure 3.**
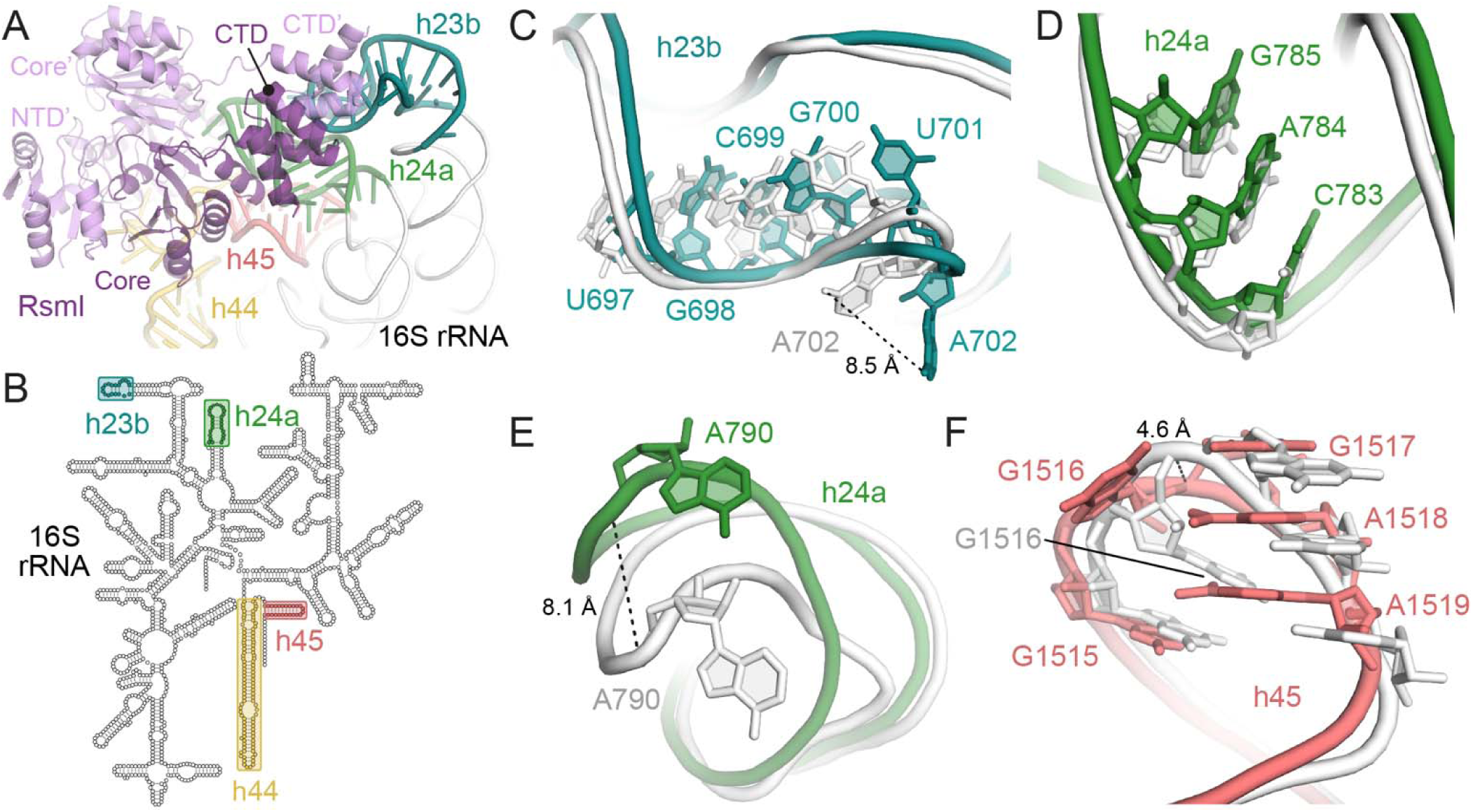
RsmI binds to a 16S rRNA surface only present in the assembled 30S subunit. (*A*) View of RsmI in contact with the four helices comprising the 16S rRNA binding surface–h44 (yellow), h45 (red), h24a (green), and h23b (cyan). (*B*) 2D representation of the 16S rRNA secondary structure, highlighting the disparate locations of the helices that form the RsmI binding surface. Comparisons between RsmI-bound 30S subunit (colored as in panel A) and free 30S (white) at the points of contact between RsmI and (*C*) h23b, (*D*) h24a stem, (*E*) h24a loop, and (*F*) h45.

At the points of contact between RsmI and the 30S subunit platform, the rRNA surface features are structurally disrupted to varying extents by enzyme binding (**Fig. 3C-F**). At h23b, RsmI contacts nucleotides U697-A702, with minimal distortion of the RNA structure with the exception of A702, where the phosphate backbone shifts and repositions the nucleobase outward from its normal position (**Fig. 3C**). RsmI engages h24a at two separate contact points: with one strand of the helical region (nucleotides C783-G785) and with the loop at the tip of the helix (nucleotide A790) (**Fig. 3D,E**). At C783-G785 there is almost no distortion of the rRNA structure, suggesting that RsmI recognizes the h24a helical structure as a rigid unit (**Fig. 3D**). In contrast, at the tip of the helix, RsmI interaction promotes a large shift of the phosphate backbone of A790 (**Fig. 3E**). Lastly at h45, RsmI contacts nucleotides G1515-G1517, inducing a smaller shift in the phosphate backbone of G1517, and causing its nucleobase to rotate slightly inward toward the helix, while the position of G1515 is mostly unaffected (**Fig. 3F**). The bases of nucleotides A1518 and A1519, also rotate inward to maintain their stacking interaction with G1517. These smaller reorganizations of h45 are facilitated by the significant movement of G1516, whose nucleobase is flipped from its canonical position on the interior of the loop to an outward facing position alongside that of G1517 (**Fig. 3F**). Collectively, the distortions RsmI imposes on its rRNA binding surface appear much more widespread and substantial than has been observed to date for other 16S rRNA methyltransferases (18–20).

### RsmI interactions with h24a and h45 are important for C1402 methylation

In contrast to interactions with h44 which are made exclusively by the RsmI catalytic protomer, residues from both protomers of the RsmI dimer mediate the contacts with the other three rRNA helices (**Fig. 4A**). At h23b, RsmI engages nucleotides U697-A702 via CTD residues of its auxiliary protomer (CTD’; **Fig. 4B**; discussed further below). At the tip of h24a, RsmI surrounds nucleotide A790 with the partially conserved residues Y103 and H104 as well as residue D137 from its catalytic and auxiliary protomers, respectively (**Fig. 4C**; **Supplementary Fig. S5C** and **Table S2**). This interaction network aligns the base edge of A790 with both the polar and hydrophobic groups of the modified amine group of C1402, in an enzyme-induced RNA-to-RNA interaction with the target nucleotide (**Fig. 4C**). Functional disruption of the A790 network requires alteration of at least two of the three RsmI residue contacts and results in only a modest decrease in RsmI activity (∼30-40%; **Fig. 4E**). This suggests that the repositioning of A790 is an important but not essential component of substrate recognition by RsmI.

**Figure 4.**
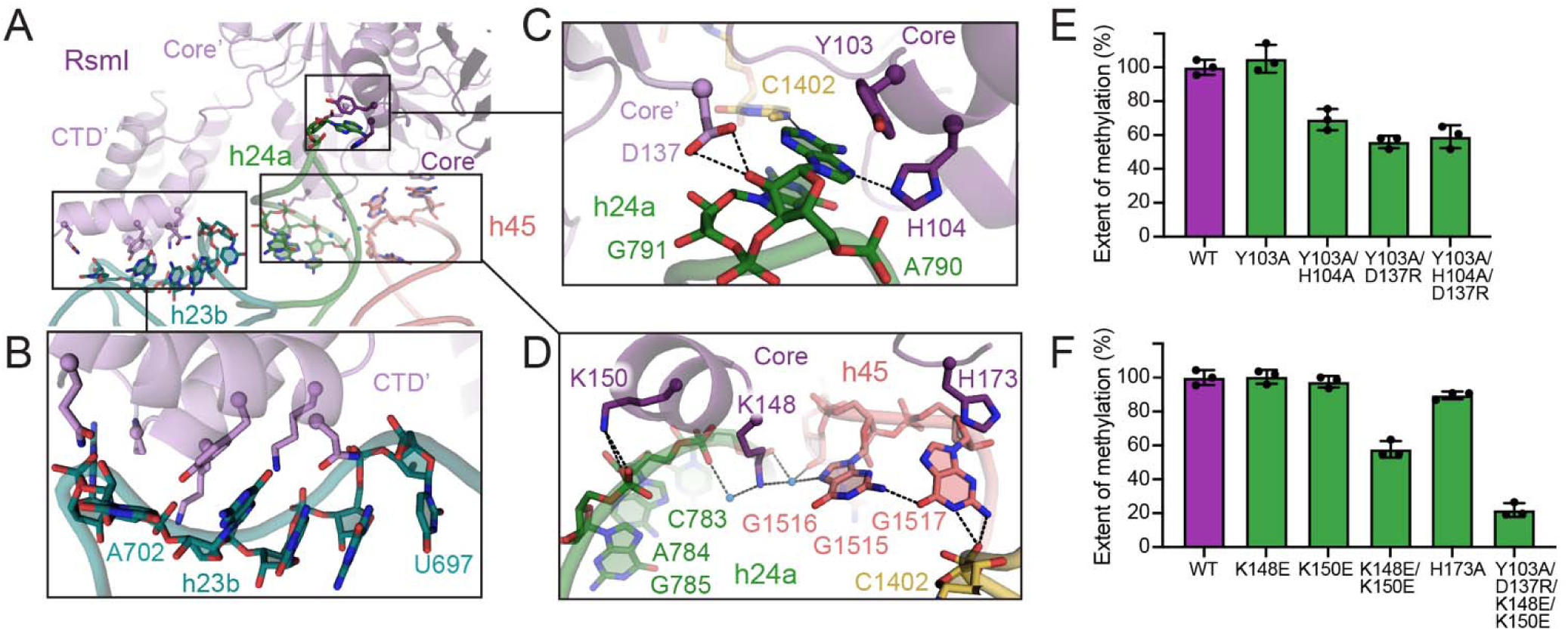
Functionally important contacts with the 16S rRNA surface comprising h23b, h24a and h45 are mediated by both RsmI protomers. (*A*) Overview of the contact points between RsmI and helices h23b, h24a and h45. Zoomed views are shown for (*B*) RsmI interaction with h23b nucleotides U697-A702, mediated by the auxiliary protomer, (*C*) the A790 interaction network, and (*D*) h24a-h45 interface network. (*E*, *F*) Methyltransferase activity assays with variant enzymes to test the functional importance of the interaction networks shown in *panels C* and *D*, respectively.

The second point of contact between RsmI and h24a (nucleotides C783-G785) and its interaction with h45 (nucleotides G1515-G1517) are mediated by an extended interaction network with the highly conserved RsmI residue K148 at its center (**Fig. 4D**; **Supplementary Fig. S5C** and **Table S2**). The water-mediated interactions made by K148 at the h24a/h45 interface result in flipping of the G1516 base to interact with G1517. G1517, in turn, is thereby positioned to hydrogen bond via its Watson-Crick base edge with the phosphate backbone of C1402, in another example of enzyme-induced RNA-to-RNA contact with the target nucleotide (**Fig. 4D**). RsmI residues K150 and H173 contribute additional interactions to this network via contacts with h24a and h45, respectively (**Fig. 4D**). While individual substitution of the RsmI residues involved in this extended network do not reduce activity, a double substitution K148E/K150E resulted in a ∼40% reduction in activity (**Fig. 4F**). Thus, as for the RsmI interaction network surrounding A790, the interactions made at the h24a-h45 interface play an important but not essential role in positioning C1402 for methylation.

Elimination of both interaction networks in a RsmI Y103A/D137R/K148E/K150E variant led to a more drastic (∼80%) reduction in activity (**Fig. 4F**), indicating that their contributions to positioning C1402 for methylation are additive. RsmI thus forms functionally important contacts with h24a and h45 through two distinct interaction networks, the combined loss of which significantly impedes the ability of RsmI to methylate C1402. As these contacts points are distant from h44, they likely play important roles in facilitating 30S subunit recognition by RsmI as well as by directly positioning C1402 through the enzyme-induced RNA-to-RNA contacts they create with the target nucleotide.

### The RsmI CTD makes functionally essential interactions with h23b

Interaction of RsmI with h23b is exclusively mediated by the previously structurally uncharacterized CTD formed by the C-terminal 42 amino acids of the enzyme. The CTD location was clearly visible in the map for both protomers, with that of the auxiliary protomer (CTD’) in direct contact with h23b (**Fig. 5A**). As this domain was missing from the published RsmI crystal structure (12) used in our initial modelling, we used AlphaFold (21) to create a preliminary structural model for the CTD, which was incorporated into the full structure for both protomers of the RsmI dimer and further refined. The newly modelled CTD consists of three α-helices (CTD-α_1_, CTD-α_2_, and CTD-α_3_) linked to the central domain of RsmI through a short flexible linker. CTD-α_3_ (residues 274-286) of the auxiliary protomer is primarily responsible for interaction of RsmI with h23b residues U697-A702. The two other helices of both protomers, CTD-α_1_ (residues 244-256) and CTD-α_2_ (residues 259-270), form an inter-protomer interface (**Fig. 5A**), with the catalytic protomer CTD-α_1_ facing away from the 30S subunit and solvent exposed.

**Figure 5.**
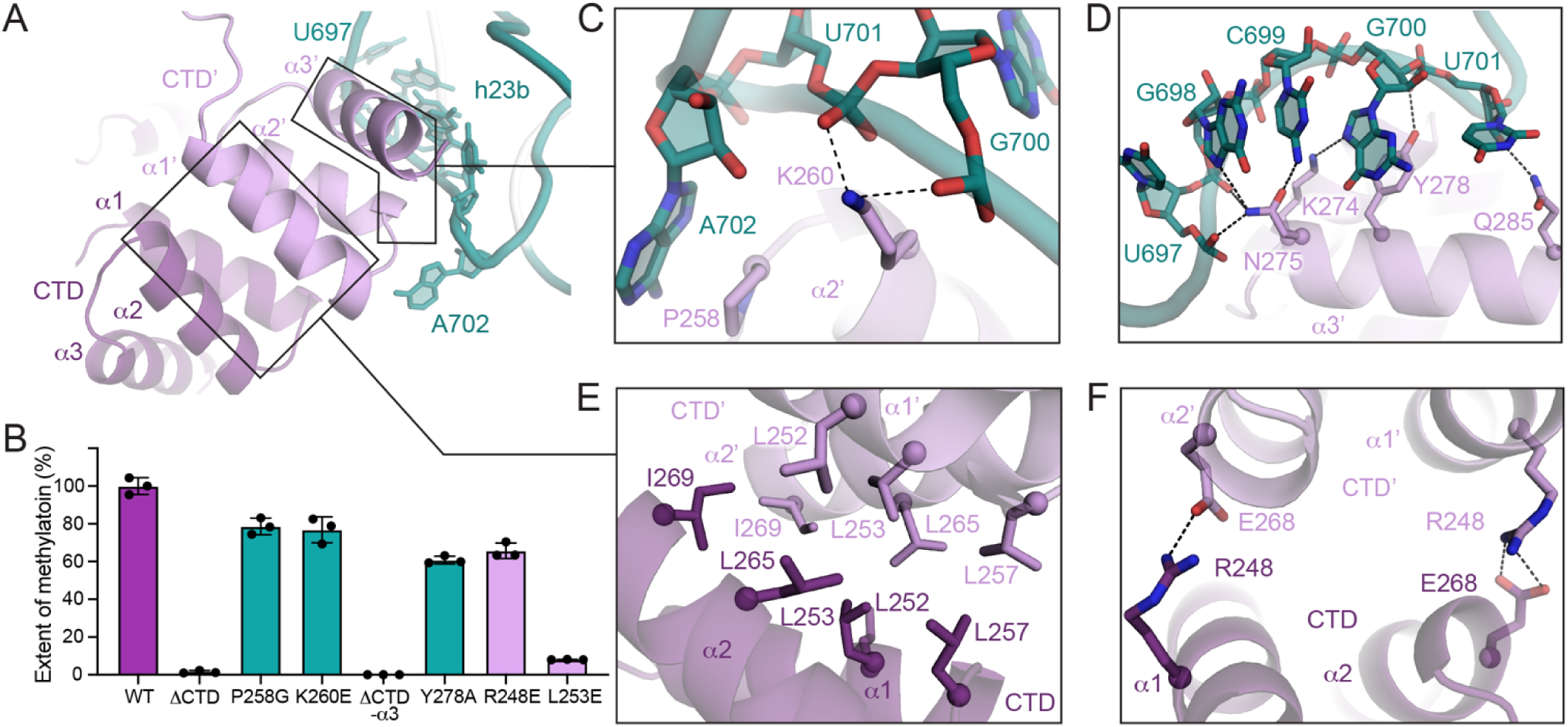
The RsmI CTD interaction with h23b is essential for C1402 modification. (*A*) Structure of the CTD dimer, consisting of three α helices in each protomer: CTD-α_1_ and CTD-α_2_ primarily engage in interprotomer-contacts and CTD-α_3_’ of the auxiliary protomer interacts with h23b. (*B*) Analysis of deletion and single amino acid variants of the RsmI CTD. Zoomed views of the interactions between h23b and (*C*) P258 and K260, and (*D*) CTD-α_3_’ residues K274, N275, Y278, and Q285. Zoomed views of the inter-protomer interaction via (*E*) a leucine-rich hydrophobic patch and (*F*) R248-mediated electrostatic contacts.

Previous structural and functional studies of RsmI lacking the CTD showed that the truncated enzyme can bind SAM and dimerize, but its ability to modify C1402 was not tested (12). Guided by our new structural insight, we examined the role of the CTD by creating a truncated RsmI variant lacking the last 50 amino acids (RsmI-ΔCTD). Remarkably, this truncated construct displayed virtually no ability to methylate C1402 (**Fig. 5B**), despite still containing all residues needed to manipulate h44 and interact with h24a and h45. The interactions made with h23b by CTD’ of the auxiliary RsmI protomer thus appear crucial for RsmI to recognize its substrate for C1402 methylation.

Two main groups of residues contribute to the interaction between the auxiliary protomer and h23b: P258 and K260 in the loop between CTD-α_1_’ and CTD-α_2_’ and at the start of CTD-α_2_’, respectively, and K274, N275, Y278, and Q285 of CTD-α_3_’ (**Fig. 5C,D**). Notably, while both P258 and K260 are highly conserved among the Gammaproteobacteria (**Supplementary Fig. S5E** and **Table S2**), individual substitution of P258 with glycine (P258G) or K260 with glutamic acid (K260E), to remove their respective stacking and electrostatic contacts results in only a small reduction (∼20%) in RsmI activity (**Fig. 5B**). As such, these interactions do not appear individually critical for h23b interaction.

We similarly evaluated the role of the CTD-α_3_ residues K274, N275, Y278, and Q285 which engage in a network of electrostatic, hydrogen bonding, and stacking interactions with the nucleobases and backbone of h23b nucleotides U697-U701 (**Fig. 5D**). The collective importance of this network was tested by constructing a truncated RsmI variant lacking CTD-α_3_ (RsmI-ΔCTD-α_3_). As for the RsmI-ΔCTD variant, the RsmI-Δα_3_ displays almost no methylation activity (**Fig. 5B**), confirming that CTD-α_3_’ is the component primarily responsible for h23b binding. Although none of the four CTD-α_3_ residues is fully conserved across all bacteria, Y278 has the highest sequence conservation (at 92% identity) and the other sites are typically at least partially conserved in their amino acid physicochemical property (**Supplementary Fig. S5E** and **Table S2**), suggesting that the collection of interactions with h23b is likely conserved across the enzyme family. Consistent with its higher sequence conservation and the observed stacking and hydrogen bonding interactions with G700, Y278 substitution with alanine (Y278A) resulted in a ∼40% reduction in methylation activity (**Fig 5B**).

Although the CTD is not necessary for RsmI dimer formation (12), given its newly identified importance for RsmI activity we next asked whether the inter-protomer contacts mediated by CTD-α_1_ and CTD-α_2_ of each protomer might potentially assist in positioning CTD-α_3_’ of the auxiliary protomer for interaction with h23b. CTD-α_1_and CTD-α_2_ of each RsmI protomer in the dimeric structure interact via a hydrophobic interface, primarily consisting of highly conserved leucine residues (**Fig. 5E**; **Supplementary Fig. S5E** and **Table S2**). Flanking this hydrophobic interface, residue R248 in each RsmI protomer forms electrostatic interactions with the side chain of E269 (**Fig. 5F**). We therefore first substituted R248 to the oppositely charged glutamic acid (R248E) to introduce charge repulsion at two contact points within the inter-protomer binding interface, which lead to a ∼40% reduction in methylation activity (**Fig. 5B**). We then substituted L253, the most conserved leucine within the hydrophobic binding interface (**Supplementary Fig. S5E** and **Table S2**), with glutamic acid (L253E) to introduce two negative charges in close proximity in the center of the hydrophobic interface, leading to a ∼90% reduction in methylation (**Fig. 5B**).

Together, these results reveal the critical role of the RsmI CTD in 30S subunit recognition. The auxiliary protomer forms essential interactions with h23b distant from the modification site in h44, mediated primarily through CTD-α_3_’. The CTD of the catalytic protomer also appears important as inter-protomer CTD contact mediated by CTD-α_1_and CTD-α_2_, primarily through the leucine-rich hydrophobic binding interface, appears to be necessary to optimally place CTD’ for h23b binding.

### h44 distortion and RsmI binding-induced RNA-RNA interactions position C1402 for modification via a divalent metal ion-dependent mechanism

As noted above, RsmI binding dramatically disrupts the h44 structure to expose and reposition C1402 within the active site of RsmI for ribose 2’-OH modification (**Figs. 2** and **6A**). This movement of C1402 is further supported by RsmI binding-induced distortions of h24a and h45 which result in new RNA-RNA interactions with the target nucleotide (**Figs. 4** and **6B,C**). With these rRNA movements, the C1402 2’-OH is positioned for covalent attachment to NM6, which is bound within the RsmI SAM binding site in a manner almost identical to SAM (12), confirming that our structure using the SAM analog accurately reflects the enzyme-cosubstrate complex (**Fig. 6D**). However, our structure trapped in an intermediate post-catalytic state additionally reveals water-mediated interactions between RsmI residues E170 and E227 with the NM6 ribose and adenine moieties, respectively, that were not previously observed in the RsmI-SAM crystal structure (12) (**Fig. 6D**).

**Figure 6.**
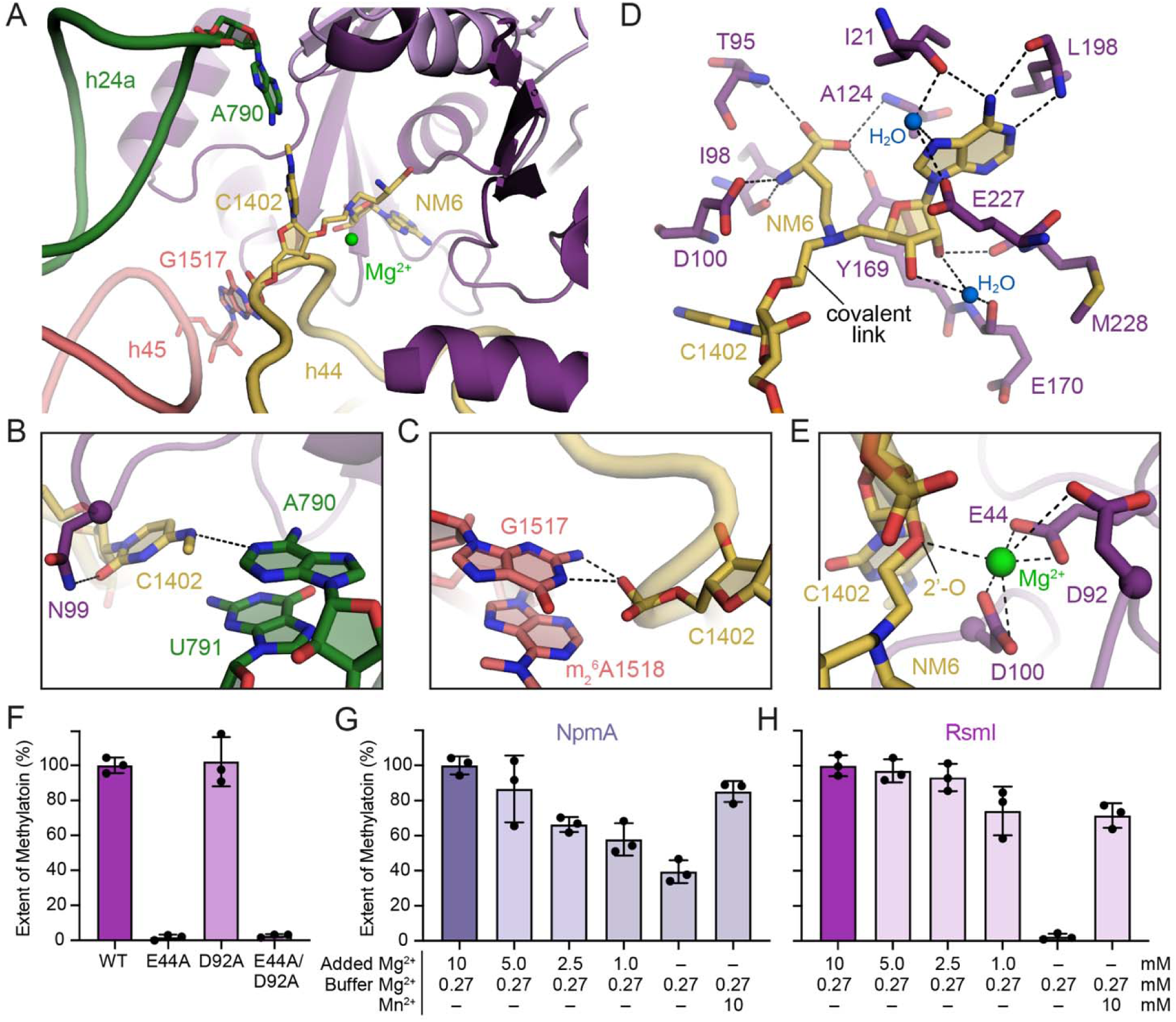
Catalysis of C1402 2’-O-methylation by RsmI depends on enzyme-induced RNA rearrangement and divalent metal ion. (*A*) Overview of target nucleotide C1402 position between RsmI, h24, and h45. Zoomed views of the RsmI-induced RNA-to-RNA interactions between C1402 and (*B*) h24a nucleotide A790 and (*C*) h45 nucleotide G1517. (*D*) View of C1402 covalently bound to NM6, in the RsmI SAM-binding pocket. (*E*) View of the Mg^2+^ coordination network between the C1402 2’-oxygen, C1403 backbone, and RsmI residues E44, D92, and D100. (*F*) Functional studies of the RsmI residues coordinating the Mg^2+^. Results of functional studies for (*G*) NpmA and (*H*) RsmI in which enzyme activity was tested with varying Mg^2+^ concentration or replacement of Mg^2+^ with Mn^2+^ in the reaction buffer.

Interestingly, our structure also identified a Mg^2+^ ion coordinated by the modified C1402 2’ oxygen, C1403 phosphate backbone, and the negatively charged RsmI residues E44, D92, and D100, which are 90-100% conserved among all RsmI enzymes (**Fig. 6E; Supplementary Fig. S4D** and **Table S2**). The position of this Mg^2+^ at the modification site suggests that it might play an important role in stabilizing the ribose 2’-OH and priming it to accept the methyl group. To test this idea, we first substituted residues E44 and D92 with alanine individually (E44A and D92A) and together (E44A/D92A). D100 was not tested as this residue was previously substituted with alanine resulting in complete disruption of SAM binding (12) and thus methyltransferase activity. While the D92A substitution did not cause a reduction in RsmI activity, the E44A and the E44A/D92A variants displayed almost complete loss of RsmI activity (**Fig. 6F**). To ensure that the lack of activity in the E44A variant does not result from loss of SAM binding, we measured the SAM affinity for wild-type and E44A RsmI using isothermal titration calorimetry (**Supplementary Fig. S7**). The K_d_ obtained for the wild-type protein (7.2 μM) was consistent with the previously reported value (12) and no reduction in binding was observed for the E44A variant (6.2 μM). These results thus directly implicate E44 in an essential role in Mg^2+^ coordination and as a residue critical for RsmI function.

To further probe the role of Mg^2+^ in the RsmI catalytic mechanism, we tested the activity of wild-type RsmI under varying Mg^2+^ concentrations, reducing from 10 mM used in the prior assays to the lowest concentration of 0.27 mM (corresponding to the divalent ion contributed to the reaction by the 30S subunit storage buffer). Notably, this Mg^2+^ concentration is similar to that used to split ribosomal subunits during their purification (22). Therefore, we expected the 30S subunit to retain its overall structural integrity at the lowest Mg^2+^ concentration but recognized the potential for a general destabilization of the RNA structure across the 16S rRNA surface necessary for RsmI interaction. Therefore, as a control, we performed equivalent methyltransferase assays with the 16S rRNA (m^1^A1408) methyltransferase NpmA, which also requires an assembled 30S subunit to function and docks on a similar rRNA surface but only minimally disrupts its structure (18). NpmA activity declines with each reduction in Mg^2+^ concentration, consistent with this enzyme docking rigidly on a pre-formed rRNA surface which becomes increasingly less stable as the divalent ion concentration is reduced. Importantly, however, even at the lowest concentration, NpmA retains about 40% activity (**Fig. 6G**). In contrast, the activity of RsmI is essentially unaffected by the reduced divalent ion concentration until a critical threshold between 1 and 0.27 mM, at which point activity is fully ablated, confirming the dependence of RsmI on the divalent ion (**Fig. 6H**). We draw two conclusions from this finding. First, RsmI activity is less sensitive to the intermediate Mg^2+^ concentrations and associated destabilization of the rRNA surface as, unlike NpmA, RsmI must distort this structure to access its target. Second, the subsequent complete loss of activity at <1 mM Mg^2+^ directly implicates this coordinated divalent metal in the RsmI catalytic mechanism. Additionally, this dependence is not specific to Mg^2+^, as replacement of Mg^2+^ with Mn^2+^ at 10 mM resulted in only 20-30% loss of activity for NpmA and RsmI (**Fig. 6G, H**).

Finally, to further probe the role of the Mg^2+^ ion in RsmI activity, we replaced the divalent ion in the reaction buffer with the hexahydrated Mg^2+^ mimic Co(NH) ^3+^, which can replicate the electrostatic function of Mg^2+^ but cannot engage in proton transfer (23, 24). For NpmA, titration of Co(NH_3_)_6_^3+^ into methylation reactions with 10 mM Mg^2+^ did not appreciably alter activity, and full activity was restored at all Co(NH_3_)_6_^3+^ concentrations in the absence of added Mg^2+^ (**Supplementary Fig. S8A**,**B**). These observations are consistent with the ability of Co(NH_3_)_6_^3+^ to replace the structural role of Mg^2+^ in proper 16S rRNA folding. In contrast, for RsmI addition of Co(NH_3_)_6_^3+^ at either Mg^2+^ concentration resulted in substantially increased RsmI activity (**Supplementary Fig. S8C**,**D**). These results suggest that RsmI requires Mg^2+^ for an essential role in coordinating and precisely positioning the C1402 2’-OH for modification, rather than a direct involvement in proton transfer, which Co(NH_3_)_6_^3+^ could not support. However, further studies are needed to determine the specific details of the contribution of Mg^2+^ to the RsmI catalytic mechanism and the basis of the unexpected observation that RsmI activity is higher in the presence of Co(NH_3_)_6_^3+^.

## Discussion

The bacterial ribosome contains ∼36 rRNA modifications primarily clustered in the functional centers of both subunits (25). The 30S subunit contains 11 of these modified rRNA nucleotides, including 9 base N-methylations, one dual base N- and ribose 2’-O-methylation (C1402), and one pseudouridine, incorporated by 11 unique enzymes (25). While much is known about the effects of these modifications on 30S structure and function (2–6), there is still a relative dearth of information about how these modifications are incorporated, in particular with respect to the timing of 30S biogenesis. The 30S P site and surrounding rRNA is heavily modified, with two nucleotides in h31 modified by the intrinsic methyltransferases RsmB (m^5^C967) and RsmD (m^2^G966), as well as three h44 nucleotides modified by RsmE (m^3^U1498), RsmF (m^5^C1407), RsmH (m^4^C1402) and RsmI (Cm1402) (7, 26–28). The enzymes that incorporate the cluster of modifications in h44 are ubiquitously conserved, yet non-essential under typical laboratory growth conditions, with their respective rRNA modifications being implicated in tuning translation (7, 29) and antibiotic susceptibility (6, 7, 29–32). While these enzymes likely function late in 30S subunit biogenesis given their shared substrate requirement of a mature or near mature 30S subunit (7, 27, 33), how they access and modify their target nucleotides, and what role(s) these methyltransferases and/or their modifications play in 30S subunit assembly have remained unclear.

To address such questions for the 2’-O-methyltransferase RsmI, we determined its mechanism of 30S subunit substrate recognition and modification using single-particle cryo-EM and complementary functional analyses. These studies revealed that RsmI accesses C1402 within the context of the assembled 30S subunit by inducing substantial distortion and unwinding of h44 via interactions mediated by its catalytic protomer. We also show that the requirement for an assembled 30S subunit as substrate arises from RsmI’s interaction with an extended 16S rRNA surface comprising h23b, h24a, h44, and h45. The interactions made by RsmI with h24a and h45 promote novel RNA-to-RNA contacts with the C1402 target nucleotide that are important for activity. Meanwhile, the functionally essential interaction with h23b, most distant from C1402, is mediated exclusively by the newly resolved CTD of the auxiliary RsmI protomer, additionally revealing why RsmI functions as a dimeric enzyme. The extensive interactions made by RsmI and the resulting distortions of the 16S rRNA surface collectively reposition the ribose 2’-OH of C1402 to receive a methyl group from the SAM co-substrate in a divalent metal-ion dependent process.

The unraveling of h44 by RsmI represents the most severe distortion of the 16S rRNA structure by a 30S subunit-targeting rRNA methyltransferase observed to date (18–20). How RsmI is able to induce such a major structural distortion in h44, including disruption of multiple base pairs in the absence of energy-driven helicase activity, can be rationalized by our structure-guided functional analyses of the key interaction clusters at h44. These data suggest that RsmI begins the process of unravelling h44 by first sequestering A1408, several nucleotides distant from the site of modification, and then using an array of conserved positive and aromatic residues to unpair the RNA duplex of h44 in a 3’ to 5’ direction, culminating in a ∼26 Å movement of C1402 from its normal position within h44 into the RsmI catalytic center. A1408 is not involved in a canonical Watson-Crick base pair within h44 as it is located opposite two dynamic nucleotides, A1492 and A1493, that unstack from the helix interior as part of their essential role in probing mRNA-tRNA interaction during decoding (34). This increases the mobility of the A1408 nucleobase, a feature that has been proposed to be exploited by the aminoglycoside-resistance m^1^A1408 methyltransferase NpmA which rigidly docks on the 30S subunit to capture A1408 between two conserved Trp residues when flipped from h44 (18). Unlike RsmI, the movement of A1408 is the only disruption of the 16S rRNA structure observed when NpmA is bound to its 30S substrate. Our data suggest that RsmI also exploits this dynamic nature of A1408 to gain an initial anchor point for h44 unwinding as blocking the movement of A1408, whether the 30S subunit has been activated or not, eliminates modification of C1402 (**Fig. 2G**). Further, supporting the idea of sequential unwinding from A1408 to C1402, the interaction networks that reposition nucleotides between these sites are also important for efficient modification, whereas interactions observed to the 5’ side of C1402 appear to be essentially dispensable.

The requirement of a structurally conserved 16S rRNA binding surface on the 30S subunit is a feature RsmI shares with several other 16S rRNA methyltransferases studied to date, including NpmA (19), the aminoglycoside-resistance m^7^G1405 methyltransferase RmtC (19), and the universally conserved intrinsic m_2_^6,6^A1518/ m_2_^6,6^A1519 dimethyltransferase RsmA (formerly KsgA) (20). However, RsmI appears distinct in the extent of its 16S rRNA interaction surface, with contacts made to h44 and h23b that span a distance of >70 Å and bury an interface of over 3000 Å^2^, more than 2.5× that buried by NpmA (18). Recognition of the distant h23b is essential for RsmI to modify C1402 as loss of these contacts made by the auxiliary protomer fully eliminate enzyme activity. A further distinction is that the three other 16S rRNA methyltransferases appear to use the 16S rRNA surface as rigid docking platforms, with rRNA structural deformations restricted to regions immediately surrounding their target sites in h44 (NpmA and RmtC) or h45 (RsmA) (18–20). In contrast, in addition to major disruption of h44, RsmI engagement with h24a and h45 results in additional distortions of the rRNA more distant from the target site which promote new enzyme-induced RNA-to-RNA contacts with the target nucleotide, potentially assisting in stabilizing C1402 for modification. Thus, RsmI appears to engage and manipulate the 16S rRNA surface in three distinct ways to achieve specific substrate recognition and access C1402 for modification: rigid docking at a distant conserved structural feature (h23b), extensive disruption of h44, and distortion of other 16S rRNA helices (h24a and h45) to assist in positioning C1402. Whether the other intrinsic 16S rRNA methyltransferases acting on h44 (RsmE, RsmF and RsmH) employ similar molecular strategies to access their targets will require corresponding studies of their 30S-enzyme complexes. Given its shared target nucleotide with RsmI, the 30S-RsmH complex would be particularly informative on the potential diversity in molecular recognition strategies used by these enzymes.

This work also revealed that RsmI unexpectedly depends on a divalent metal ion for catalysis of C1402 ribose 2’OH methylation. The well-studied and essential bacterial m^1^G37 tRNA methyltransferase TrmD similarly requires a divalent metal ion which is thought to increase the nucleophilicity of the N1 position and stabilize the developing negative charge on O6 during methyl transfer (35). Like TrmD, RsmI surrounds the divalent metal ion with a collection of negatively charged amino acids to help chelate the ion. However, in contrast to TrmD, where metal coordination is completed by the carboxylic tail of SAM and, presumably, by the nitrogen base of its substrate G37 (36), metal coordination in the RsmI-30S complex is completed by the C1403 phosphate backbone and C1402 2’-OH group without interaction with the cosubstrate. While to our knowledge RsmI represents the first characterized example of a metal-dependent RNA ribose 2’-OH methyltransferase, direct involvement of the target oxygen atom in metal coordination may be a feature shared across other metal-dependent O-methyltransferases as it is also present among metal-dependent catechol and flavonoid O-methyltransferases (37–39). For example, RsmI and the alfalfa caffeoyl coenzyme A 3-*O*-methyltransferase share strikingly similar metal coordination arrangements despite their distinct substrates. In this enzyme, the metal ion is proposed to deprotonate its target hydroxyl group and maintain the resulting oxyanion near SAM’s reactive methyl group (38). However, our studies suggest that the RsmI-bound metal ion likely does not engage in direct proton transfer but is instead coordinated by conserved and functionally critical acidic residues to precisely position the ribose 2’-OH for modification. As both an RNA-modifying and metal-dependent O-methyltransferase, RsmI may represent the first of a unique subclass of SAM-dependent RNA methyltransferases.

The timing of rRNA modification incorporation during 30S biogenesis remains an area of active investigation. For some 16S rRNA modifying enzymes, their ability to access their target nucleotide depends on prior folding of 16S rRNA and/or binding of ribosomal proteins. For instance, the binding of ribosomal proteins S7 and S19 to 16S rRNA results in an rRNA conformational switch during 30S subunit assembly which acts as a barrier separating the timing of h31 modifications incorporated by RsmB and RsmD which occur before and after S7/S19 binding, respectively (28). On the other hand, the action of other 16S rRNA modification enzymes of the bacterial small subunit have been proposed to serve as quality control check points in 30S subunit assembly, dictating progression to later stages of the process. For example, binding and 16S rRNA modification by RsmA has been suggested to act as a final checkpoint for the integrity of 30S platform assembly while also allowing time for completion of other late-stage 30S assembly processes (2, 20, 40). The action of RsmI late in 30S assembly and the way it engages and manipulates the folded 16S rRNA suggests the potential for a similar role in 30S subunit biogenesis and quality control. For instance, coordinated recognition of distant sites (h44 and h23b) implicates RsmI as quality control factor for both the correct folding of the 30S platform and the relative placement of h44, the absence of which could stall RsmI action.

Additionally, the extensive unfolding of h44 comprising a key functional region of the ribosome offers a potential “reset button” and opportunity for correct refolding of the ribosomal P site and adjacent decoding center in the event that they fall into a kinetic folding trap during assembly (41). This idea is further reinforced by our finding that RsmI can withstand destabilization of the 30S surface during C1402 modification (**Fig. 6H**). Further, given the overlapping binding sites of RsmI with either mRNA, translation initiation factor 3, or H69 of the 50S subunit in the context of a functional 70S (42, 43), the presence of RsmI on pre-mature 30S subunits could also serve to impede binding of other factors, effectively removing the 30S subunit intermediates from participating in translation. We also note that the necessity of all elements of the extreme unravelling of h44 observed *in vitro* may depend on the timing of C1402 modification by RsmI in the bacterial cell. The free mature 30S subunit predominantly adopts an inactive conformation (17), and our data show that RsmI exhibits greater activity on a mixed populated of 30S subunits that have not been activated *in vitro* (**Fig. S5**). However, the complete loss of activity observed when RsmI interactions are disrupted at the most distant h44 contact (A1408; **Fig. 2G**), regardless of whether 30S subunit is activated or not, demonstrate that h44 must be fully unfolded by RsmI, as we observe in the structure, to enable C1402 modification. There also remains the possibility that RsmI modifies a pre-mature 30S subunit, prior to proper docking of h44, which is thought to be one of the last steps in 30S assembly (44, 45). Regardless, any late-stage 30S assembly intermediate on which RsmI acts must at least be sufficiently formed such that h23b of the 30S platform and h44 are present and in position to allow simultaneous contact by RsmI. As such, we expect that the extensive unpairing of h44 base pairs to access and specifically target C1402 would be a key element of the RsmI mechanism regardless of the point in assembly that modification occurs. However, discerning the true nature of any late stage 30S intermediate(s) serving as the RsmI substrate and the extent of necessary h44 rearrangement will require further studies.

Our structure-guided approach to deciphering the RsmI mechanism of action has expanded our understanding of the breadth of rRNA methyltransferase activity in terms of 30S subunit recognition, rRNA manipulation and metal ion dependence in rRNA modification. Moreover, our studies provide further support for the concept of intrinsic bacterial methyltransferases as ribosome assembly checkpoints, helping to contextualize how we view such intrinsic bacterial methyltransferases beyond just depositors of methyl modifications.

## Materials and Methods

### Generation of an E. coli ΔrsmI strain for C1402 hypomethylated 30S subunit isolation

An *E. coli* strain lacking RsmI activity was generated by replacing *rsmI* with the chloramphenicol acetyltransferase gene (*cat*) using conventional gene replacement strategies (7, 46) (**Supplementary Fig. S1A**). First, *cat* and its promoter were amplified from the plasmid pZS21 (47, 48) using primers to incorporate 40 nucleotide extensions homologous to *rsmI* at each end (**Supplementary Table S3**). The resulting *cat* cassette was used to transform via electroporation *E. coli* strain DY378, which endogenously expresses the lambda red homologous recombination machinery under temperature control (49). Finally, P1 phage transduction (50), was used to transfer the Δ*rsmI* genotype to *E. coli* BW25113 (46). Replacement of *rsmI* with *cat* was verified through PCR amplification of genomic DNA extracted using the DNeasy Blood and Tissue Kit (Qiagen) (**Supplementary Fig. S1B** and **Table S3**). Wild-type BW25113 and 30S-Δ*rsmI* ribosomal subunits were prepared by iterative rounds of high-speed centrifugation to isolate the ribosomal fractions of the cell followed by sucrose gradient centrifugation at low Mg^2+^ concentration to separate the 30S subunit from other ribosomal populations, as done previously (18). Activation of the 30S subunit is required for h44 and the decoding center to adopt their canonical conformation in the 30S subunit *in vitro* (15, 16). Unless explicitly stated otherwise, all 30S subunits were activated by incubation at 42 °C for 5 minutes and then room temperature for 10 minutes to ensure the consistent nature of all 30S species prior to cryo-EM specimen preparation and methyltransferase assays with RsmI variants. 30S-Δ*rsmI* was shown to be unmethylated at C1402 (and thus a substrate for RsmI) using an *in vitro* methylation assay as described further, below (**Supplementary Fig. S1C,D**).

### RsmI protein expression, purification and site-directed mutagenesis

The sequence encoding RsmI was amplified from *E. coli* BW25113 genomic DNA (**Supplementary Table S3**) and subcloned into a pET44a vector with an N-terminal hexahistidine tag (pET44-RsmI). This plasmid was used to transform *E. coli* BL21(DE3) cells with subsequent growth at 37 °C in lysogeny broth containing ampicillin (100 µg/ml). Protein expression was induced at mid-log phase (∼0.6 OD_260_) with 1 mM isopropyl β-D-1-thiogalactopyranoside and growth continued for another 3 hours before being harvested via centrifugation at 4°C. Cells were resuspended in lysis buffer (50 mM HEPES-NaOH, pH 7.6, 1 M NaCl, 10 mM imidazole, 0.5% Triton X-100, and 2 mM β-mercaptoethanol containing the protease inhibitors PMSF and benzamidine) and lysed by sonication (Misonix Sonicator 3000 with microtip: 7.5-min total sonication time, 1-s on, 1-s off, output level 5.0). Cell lysates were cleared by centrifugation at 4°C and then dialyzed three times (2 x 3 hours and overnight) against a high salt buffer (50 mM HEPES-NaOH, pH 7.6, 2 M NaCl, 10 mM imidazole, and 2 mM mM β-mercaptoethanol) to remove any SAM copurifying with RsmI. The lysate was then dialyzed for 3 hours against the same buffer but with reduced salt (500 mM NaCl) and applied to a Cytiva HisTrap FF crude 1 mL column on an ÄKTA Purifier 10 System equilibrated in the same buffer. The column was washed for 15 column volumes with the same buffer but containing 25 mM imidazole and protein subsequently eluted by increasing imidazole to 500 mM. The eluted protein sample was further purified using a Superdex75 16/60 gel filtration column (Cytiva) equilibrated in gel filtration buffer (10 mM HEPES-NaOH, pH 7.6, 300 mM NaCl, and 2 mM β-mercaptoethanol).

RsmI variants were generated in the pET44-RsmI plasmid using the MEGAWHOP mutagenesis method (51). Primers used for first round PCR (to generate the “mega” primers) are listed in **Supplementary Table S3**. All RsmI variant proteins were expressed as for the wild-type protein. A simplified purification strategy was used for wild-type and variant RsmI proteins for use in functional assays, using NEBExpress Ni Spin Column for affinity purification with minor modifications: three wash steps were done with increasing imidazole concentrations of 25, 50, and 100 mM before elution with 500 mM imidazole. Protein folding/ stability and prep-to-prep quality control was done using nano differential scanning fluorimetry on a NanoTemper Tycho NT.6.

### Cryo-EM specimen preparation and data collection

A 3 µl mixture of wild-type RsmI (1.2 µM), 30S-Δ*rsmI* (0.4 µM), and SAM analog NM6 (24 µM), prepared as previously described (13), was applied to glow-discharged C-flat 1.2/1.3 Au 50 300 mesh grids. Grids were blotted at 4°C for 5 seconds at 100% humidity and frozen in liquid ethane using a Vitrobot Mark IV (Thermofisher). Cryo-EM data (8,164 micrographs) were recorded as movies with a defocus range from −0.4 to −2.9 μm at 105,000× nominal magnification (0.41 Å/pixel) on a Titan Krios 300 kV (TEM) with a Gatan K3 direct electron detector at the National Center for CryoEM Access and Training (NCCAT). Movies were collected at a dose rate of 27.37 e^-^/Å^2^/s with a total exposure of 1.8 seconds, for an accumulated dose of 49.27 e^-^/Å^2^. Intermediate frames were recorded every 0.05 seconds for a total of 40 frames per micrograph.

All image processing steps were conducted in CryoSPARC (52) following the workflow outlined in **Supplementary Fig. S2**. Movies were dose-weighed and corrected for motion drift using Patch Motion Correction with the output micrographs down sampled by a binning factor of 0.5. The contrast transfer function (CTF) was estimated using Patch CTF Estimation, and images with a CTF fit resolution > 5 Å were discarded (**Supplementary Fig. S2A**). Automatic blob picking selected 3,781,314 particles, which then underwent particle picking followed by extraction with a box size of 448 pixels, down-sampled to 128 pixels for faster initial particle curation, culminating in an initial set of 2,140,666 particles (**Supplementary Fig. S2B**). A series of 2D classification jobs to remove non-30S junk decreased the particle count to 1,380,845 and was followed by 3D particle sorting through iterative rounds of *ab initio* reconstruction and heterogenous refinement to further improve the dataset. Through this process, the resulting 938,684 30S particles were split into six reconstructions differing only in their occupancy of RsmI (**Supplementary Fig. S2B**). The reconstruction with highest RsmI occupancy (solid line box in **Supplementary Fig. S2B**), consisting of 356,021 particles (∼38% of non-junk 30S particles), was retained for further refinement. Additionally, the reconstruction with the lowest occupancy of RsmI (288,883 particles; dashed line box in **Supplementary Fig. S2B**) was used as the starting point to generate a map for the free 30S-Δ*rsmI* after further processing (see below). The remaining 293,780 particles from the four reconstructions with intermediate RsmI occupancy were discarded (**Supplementary Fig. S4B**).

Particles with high RsmI occupancy were re-extracted at a box size of 448 pixels and underwent iterative rounds of Homogenous Refinement, Global CTF Refinement, Local CTF Refinement, and Reference Based Motion Correction to produce a final map of the complex at 2.12 Å (**Supplementary Fig. S2C**). Further particle curation through iterative rounds of 3D Classification using a mask around the enzyme and accompanied by Heterogenous Refinement resulted in a decreased particle count of 264,656 particles, which formed a consensus 2.19 Å map after Homogenous Refinement. (**Supplementary Fig. S2D**). From this map, two separate strategies were employed. First, the particle density around RsmI and its 30S binding interface was subtracted and RsmI and its 30S contact surface locally refined using masked local refinement, leading to a 2.55 Å focused map (**Supplementary Fig. S2E**). For the second strategy, remnant 30S head-induced heterogeneity was removed and RsmI occupancy further improved through successive 3D classifications at low (16 Å) and high (6 Å) filter resolution, resulting in a final particle count of 74,553 particles. These particles underwent homogenous refinement followed by masked local refinement without particle subtraction, resulting in a 2.42 Å map of the entire complex (**Supplementary Fig. S2F**). All masks were created using a combination of ChimeraX (53) and CryoSPARC (52). Local resolution estimates of both maps were generated using CryoSPARC (52) and visualized with ChimeraX (53) (**Supplementary Fig. S3**). Sharpened maps used for **Fig. 1** and **Supplementary Fig. S3** were generated using DeepEMhancer (54).

To generate a map for free 30S-Δ*rsmI* (**Supplementary Fig. S4B**), the 288,883 particles from the lowest-occupancy reconstruction after the initial 3D particle curation (dashed line box in **Supplementary Fig. S2B**) underwent further processing to remove as many particles with RsmI as possible. After additional iterative rounds of heterogenous refinement followed by successive 3D classification jobs filtered at 6 Å and 12 Å resolution, respectively, a collection of 58,000 particles remained with virtually no RsmI present. These particles were further refined following a similar strategy as described above to generate the final 2.91 Å final map for free 30S-Δ*rsmI*.

### Model building and refinement

The 2.55 Å focused map displayed better map quality for RsmI and its 16S rRNA interface, while the 2.42 Å map of the full complex displayed better quality for the rest of the 30S body. Both maps were therefore used for model building and refinement. An empty 30S subunit structure (PDB 7OE1) (55) and the RsmI NTD and core domains from the RsmI-SAM complex crystal structure (PDB 5HW4) (12) were rigid-body fitted into both maps using ChimeraX (53) to create a starting model. After initial real-space refinement with global minimization, simulated annealing, local grid search, and ADP-factor refinement against the full map in Phenix 1.21-5207 (56), the model was truncated to the regions within the focused map (both RsmI protomers and 16S rRNA nucleotides 682-708, 780-802,1399-1413,1486-1494, and 1507-1527) and further refined against the focused map using Phenix (56) (**Supplementary Table S1)**. Residues of h44 interacting with RsmI were substantially reorganized compared to their normal locations in the 30S subunit and were manually re-built in Coot 0.9.8.95 (57) with real space refinement. Additionally, Coot (57) was used to *de novo* model the NM6-modified C1402 residue and to make all other necessary adjustments of residues for optimal fit in the map. The refined truncated region was then combined with the rest of the model, which underwent additional rounds of Phenix (56) real-space refinement with global minimization and ADP refinement coupled with geometry minimization and manual model adjustment in Coot (57), using both maps to refine distinct parts of the model. Inherent movement of the 30S head domain resulted in poorer map quality for this region and limited our ability to accurately model the corresponding 16S rRNA residues and associated ribosomal proteins. Additionally, two highly flexible residues (1397 and 1398) which link h44 to the 30S head were not included in the model. Residues 1494-1506 also displayed disorder but could be modeled and refined using a low-resolution version of the full map which showed visible density for this region (**Supplementary Fig. S4E**).

A model of the 7-amino acid linker and 42-amino acid CTD was generated using the AlphaFold 3 Server (21). Two copies of this model, one for each RsmI protomer, were then rigid-body fitted into available density on the 2.55 Å focused map using ChimeraX (53) and refined using Phenix (56). The additional domains were then covalently linked to the initial RsmI (NTD-core) structure using a combination of PyMOL (58) and Coot (57). The contacts between RsmI and the 30S platform were then further refined using Phenix (56) with finer adjustments made in Coot (57), as before. Water and Mg^2+^ were added manually using Coot (57). The free 30S-Δ*rsmI* structure was refined using the KNexPHENIX workflow (59), with waters and Mg^2+^ added manually using Coot (57).

The final models were validated in Phenix (56), and the coordinates and maps deposited in the Protein Data Bank (accession codes 9PZG and 10FZ for the RsmI-30S complex and free 30S-Δ*rsmI*, respectively) and Electron Microscopy Database (accession code EMD-72071 and EMD-75144). All parameters pertaining to data collection, image processing, model building and refinement, and model validation are summarized in **Supplementary Table S1**. Structural alignments were performed and solvent exposed surface area was calculated in PyMol, the latter using the get_area command with default 1.4 Å probe radius. Figures showing maps and structural images were generated using PyMOL (58) or ChimeraX (53).

### RsmI conservation analysis

RsmI methyltransferase sequences were retrieved from UniProt using the InterPro identifier IPR018063. Sequence redundancy was removed by applying a 90% sequence identity cutoff (UniRef90) and clusters with fewer than 50 sequences were excluded. This process yielded a final set of 198 representative RsmI sequences, covering all major bacterial groups including Proteobacteria (Alpha-, Beta-, Gamma-, and others), Firmicutes, Actinobacteria, Cyanobacteria, and thermophilic bacteria. Multiple sequence alignment was performed in Geneious, and a neighbor-joining phylogenetic tree was generated to identify and isolate the Gammaproteobacterial RsmI clade. Residue propensities were calculated and sequence logos generated in Geneious software.

### [^3^H]-SAM methyltransferase assay

Methylation of 30S subunit by wild-type and variant RsmI proteins was quantified using a filter-based enzyme assay with [^3^H]-SAM. In initial assays to test RsmI activity on the 30S-Δ*rsmI* subunit, wild-type RsmI (0.4 µM) was incubated with [^3^H]-SAM (0.84 µM) and either wild-type 30S or 30S-Δ*rsmI* subunit (0.4 µM) in Buffer G (5 mM HEPES-KOH, pH 7.5, 50 mM KCl, 10 mM NH_4_Cl, 10 mM magnesium acetate, and 6 mM β-mercaptoethanol) in a final reaction volume of 10 µl. The reaction was incubated for 90 minutes at 37°C and then quenched with 90 µl of ice-cold 5% trichloroacetic acid (TCA). The quenched reaction was transferred to a glass microfiber filter, which was washed 3× with 200 µl 5% TCA and 2× with 200 µl 70% ethanol. After drying overnight, ^3^H retained on the filter (i.e., extent of 30S methylation) was determined by scintillation counting using a Beckman Coulter LS6500 liquid scintillation. A control assay with no added enzyme was used to determine background values which were then subtracted from the experimental counts.

To establish optimal conditions for comparisons between variant RsmI enzymes, a time-course experiment was conducted with wild-type RsmI (100 nM), [^3^H]-SAM (840 nM), and 30S-Δ*rsmI* (100 nM) in Buffer G in a total volume of 90 µl. The reaction was incubated for 120 minutes at 37 °C, with 10 µl aliquots removed and quenched with 90 µl 5% TCA at 0, 1, 2, 5, 10, 20, 40, 60, 90, and 120 minutes. Subsequent steps and data analysis were performed as described above (**Supplementary Fig. S1D**). A single 20-minute time point where wild-type RsmI reaches 70-80% of maximum methylation was subsequently selected for the analyses of all RsmI variants performed under otherwise identical conditions. All assays with RsmI variants were performed alongside similarly purified wild-type RsmI and with background control, and data were normalized to values for wild-type RsmI.

Time course methylation assays comparing activated and non-activated 30S-Δ*rsmI* were conducted as above, but with the aliquots quenched at 0, 2.5, 5, 10, 20, 40, and 60 minutes. Additional experiments in which 30S subunit substrate concentration was varied (25 to 400 nM) were conducted at 5 and 40-minute time points. All methylation data comparing activated and non-activated 30S-Δ*rsmI* were normalized to the highest non-activated 30S-Δ*rsmI* value.

Finally, methylation assays with varying divalent ion (Mg^2+^, Mn^2+^ or Co(NH_3_)_6_^3+^) concentrations were conducted essentially identically other than variations in Buffer G to alter the final added magnesium acetate (0 to 10 mM), MnCl_2_ (0 or 10 mM), or Co(NH_3_)_6_^3+^ concentrations (0 to 1 mM). All assays additionally contained 0.27 mM magnesium acetate from the 30S subunit storage buffer. These data were normalized to the 10 mM Mg^2+^ conditions for either RsmI or NpmA. All methylation assay data were plotted in GraphPad Prism 10

### Isothermal Titration Calorimetry

SAM binding affinities for wild-type and E44A variant RsmI proteins were measured in triplicate using an Auto-iTC_200_. Purified proteins were dialyzed exhaustively against gel filtration buffer lacking β-mercaptoethanol and the final dialysate was used to prepare the SAM solution (1 mM). Titrations were conducted at 25 °C and comprised 19 x 2 µl injections of SAM into the sample cell containing either wild-type RsmI or E44A (29-44 µM), with 120 s between injections to allow the power signal return to baseline. Raw titrations were processed (baseline and integration window adjustments, peak integration and subtraction of heats of dilution) and the resulting data integrated fit to a one site model via nonlinear regression using the modified Origin 7.0 software supplied with the instrument.

## Data Availability

Cryo-EM maps and atomic models have been deposited in the Electron Microscopy Databank (EMD-72071) and Protein Databank (PDB ID: 9PZG), respectively. All other data are included in the manuscript and/or supporting information.

## Acknowledgements

This study was supported by the National Institutes of Health awards R01 AI088025 (to G.L.C. and C.M.D.), T32 GM135060 (to M.I.B.) and F31 AI186518 (to M.I.B.). C.M.D. is a Burroughs Wellcome Fund Investigator in the Pathogenesis of Infectious Disease recipient. This study was also supported by the Robert P. Apkarian Integrated Electron Microscopy Core (IEMC) at Emory University, which is subsidized by the Emory School of Medicine and Emory College of Arts and Sciences. Some of this work was performed at the National Center for CryoEM Access and Training (NCCAT) and the Simons Electron Microscopy Center located at the New York Structural Biology Center, supported by the NIH Common Fund Transformative High Resolution Cryo-Electron Microscopy program (U24 GM129539, and NIGMS R24 GM154192) and by grants from the Simons Foundation (SF349247) and NY State Assembly. We would like to thank Dr. Marcin Grabowicz for advice on generating the Δ*rsmI* strain and providing the *E. coli* DY378 strain and pZS21 plasmid.

## Notes

### Competing Interest Statement

The authors have declared no competing interest.

### Summary of Updates

Additional data added in main figure 2 and new supplementary Fig S5, and associated Results text. Additional structure fully determined and deposited (details in updated Table S1)

