## Supplementary Information for "Mechanism of 30S subunit recognition and modification by the conserved bacterial ribosomal RNA methyltransferase RsmI"

**This PDF file includes:**

Figures S1 to S8

Tables S1 to S3

**Supplementary Figures**

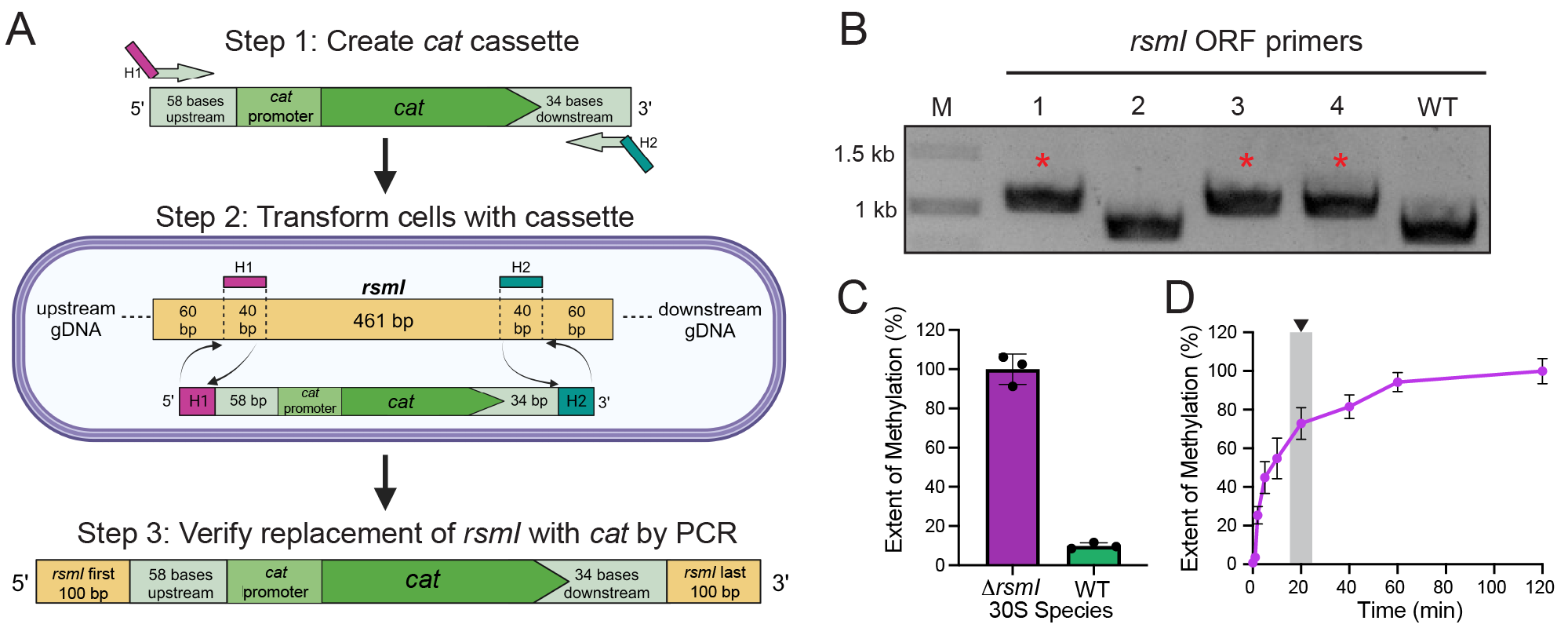

**Fig. S1.** **Sample preparation for structure-function analyses of the RsmI-30S complex.** (*A*) Scheme depicting *rsmI* deletion by replacement with a DNA cassette encoding the chloramphenicol acetyltransferase gene (*cat*) and its promoter. H1 = 5’ homology region, H2 = 3’ homology region. (*B*) Verification of *cat* replacement of *rsmI* via PCR. Colonies (1-4) were screened by PCR amplification of genomic DNA using primers targeting the intact 5’ and 3’ ends of the *rsmI* open reading frame (“*rsmI* ORF primers”). A higher molecular weight band confirms insertion of *cat* (red asterisk). WT = wild-type, M = marker. (*C*) Single time point *in vitro* methylation assay reveals that the activity of RsmI with 30S-Δ*rsmI* is significantly greater than with wildtype (WT) 30S (where this site is already modified by endogenous RsmI), indicating that 30S-Δ*rsmI* is a suitable substrate for structural studies. (*D*) Time-course methylation assay for RsmI and Δ*rsmI* 30S at a 1:1 enzyme to 30S ratio shows that RsmI begins to plateau in its activity at around 20 minutes (indicated by shaded region and arrowhead).

*[Image is shown on the following page]*

**Fig. S2.** **Cryo-EM processing workflow.** (*A*) Preliminary steps including micrograph movie import to CryoSparc, pre-processing in the form of motion correction and CTF estimation, and particle blob picking. (*B*) Particle Curation in the form of particle picking, 2D classification, and 3D sorting through ab-initio reconstruction and heterogenous refinement. (*C*) Iterative refinements jobs in the form of homogenous refinement, local and global CTF refinement, and reference-based motion correction to improve map resolution. (*D*) Additional particle curation in the form of masked 3D classification and heterogenous refinement followed by homogenous refinement to achieve a consensus volume. From this volume, two paths were chosen: (*E*) particle density around RsmI and its tertiary binding surface was subtracted to allow masked local refinement of the enzyme and its 16S rRNA interface to produce a 2.55 Å focused map, and (*F*) further 3D classification with variable filter resolutions was undertaken to remove 30S head-induced heterogeneity and further improve enzyme occupancy to produce, after additional refinement, a 2.42 Å map of the whole complex.

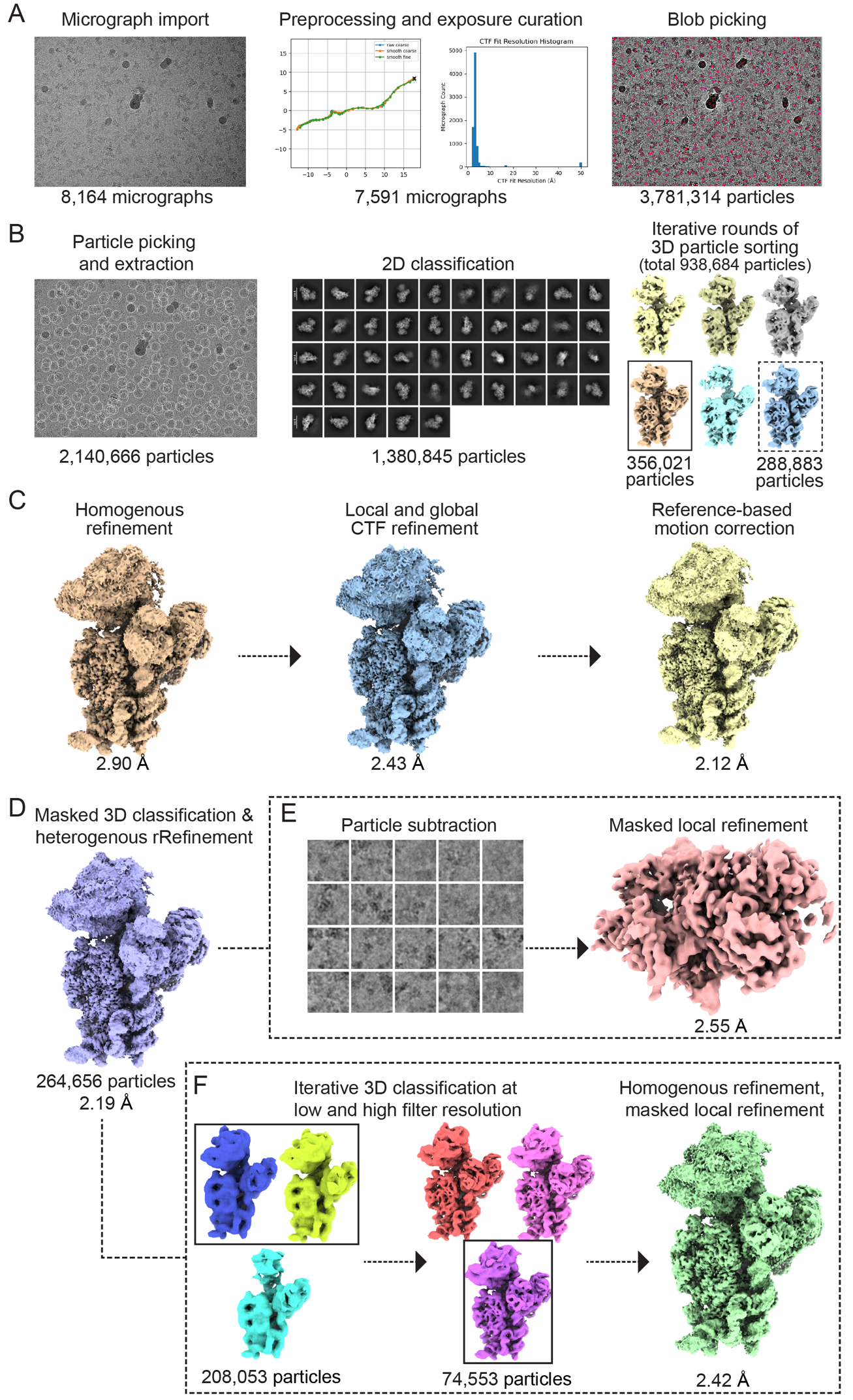

**Fig. S2.** **Cryo-EM processing workflow.** *[Full legend is provided on the previous page]*

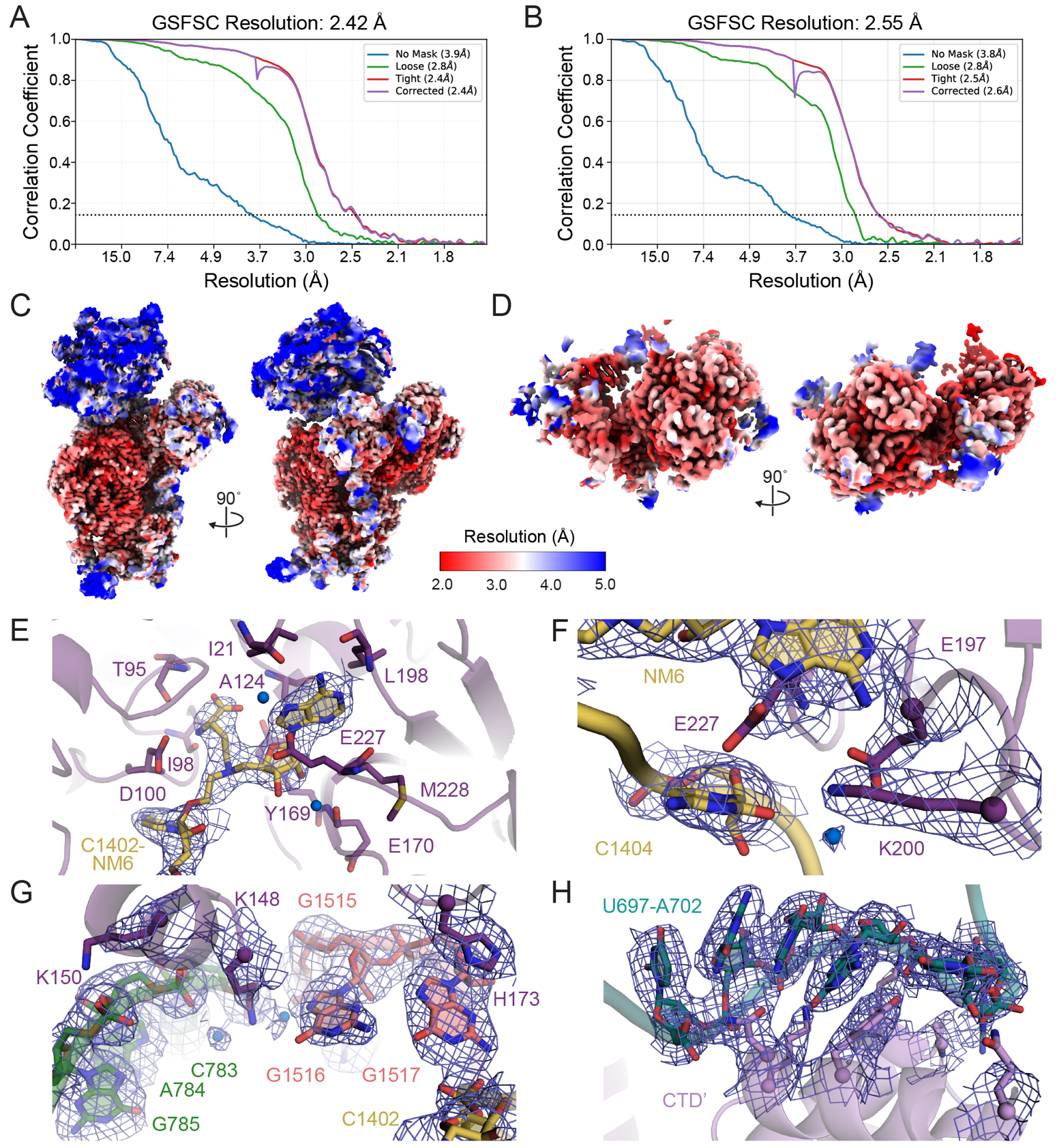

**Fig. S3. Map quality analyses.** GSFSC curves of the (*A*) 2.42 Å map of the entire RsmI-30S complex and (*B*) 2.55 Å map of RsmI and its 16S rRNA binding surface. Local resolution estimates shown as representative views for (*C*) the full RsmI-30S subunit complex and (*D*) RsmI and its 16S rRNA binding surface. Representative views of map quality for the 2.55 Å focused map highlighting (*E*) the target nucleotide C1402 covalently attached to the SAM analog NM6 (contour level 17σ) and RsmI residues in contact with (*F*) h44 nucleotide C1404 (contour level 15σ for protein/ RNA and 9σ for water), (*G*) h24a and h45 (contour level 15σ for protein/ RNA and 7σ for water), and (*H*) h23b (contour level 15σ for RNA and 10σ for RsmI).

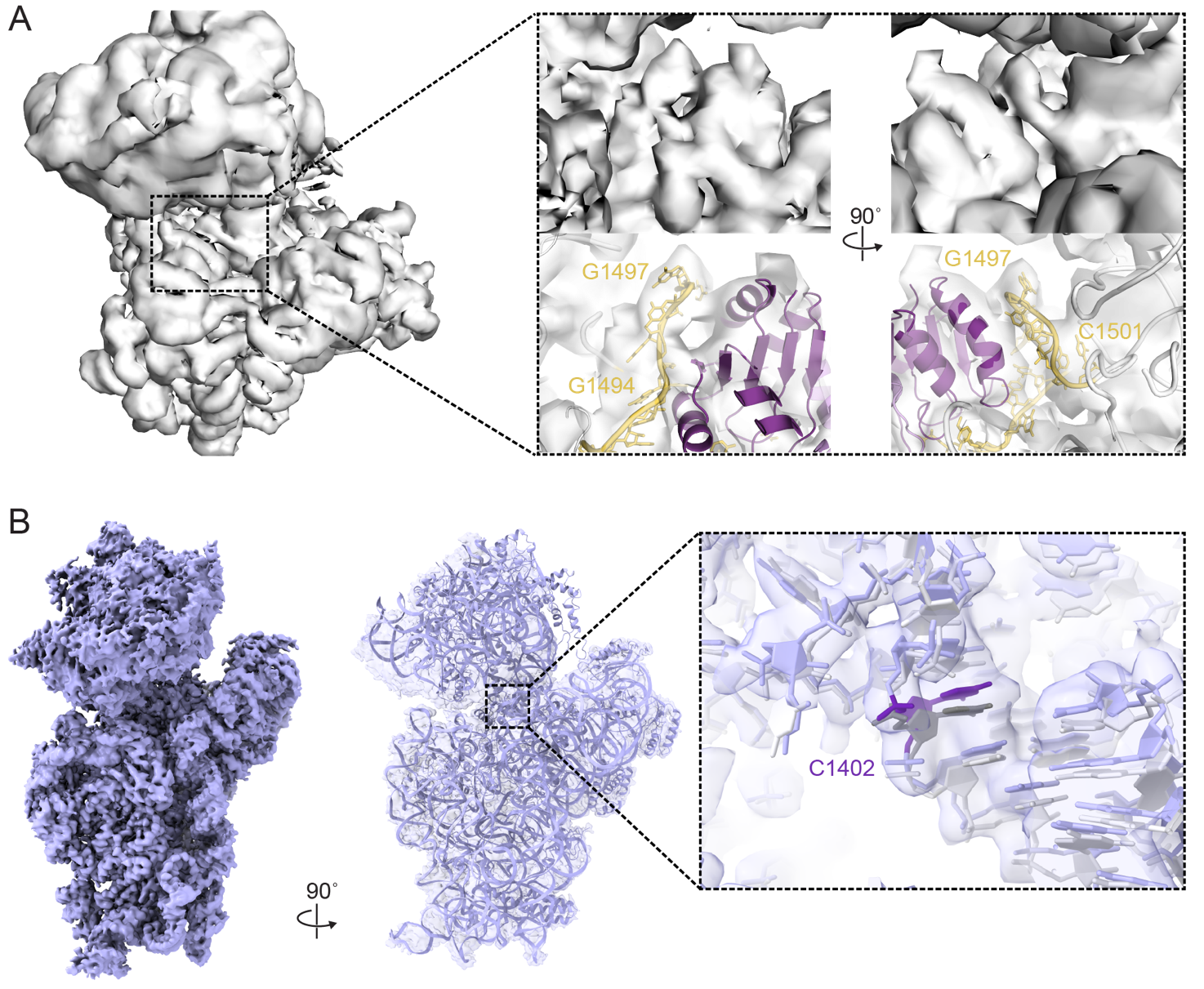

**Fig. S4. Additional maps generated during cryo-EM processing.** (*A*) Low-resolution map (6 Å) of the entire RsmI-30S complex with zoomed-in views of the map (*top*) corresponding to the dynamic h44 region (residues 1494-1501) and the final model in the semi-transparent map (*bottom*). (*B*) 2.91 Å map of the free 30S-Δ*rsmI* (*left*) with the free 30S-Δ*rsmI* structure (**Table S1**, PDB 10FZ) overlaid (*right*). The zoomed-in view shows the h44 region around C1402 (violet) from the free 30S-Δ*rsmI* structure superimposed on a wild-type 30S model (PDB 7OE1; C1402 in dark grey), with both models overlaid on the free 30S-Δ*rsmI* map (ChimeraX threshold = 0.119). The two models align well (RSMD = 0.90) and occupy the same density in the map, indicating that the free 30S-Δ*rsmI* and wild-type 30S are essentially identical in structure.

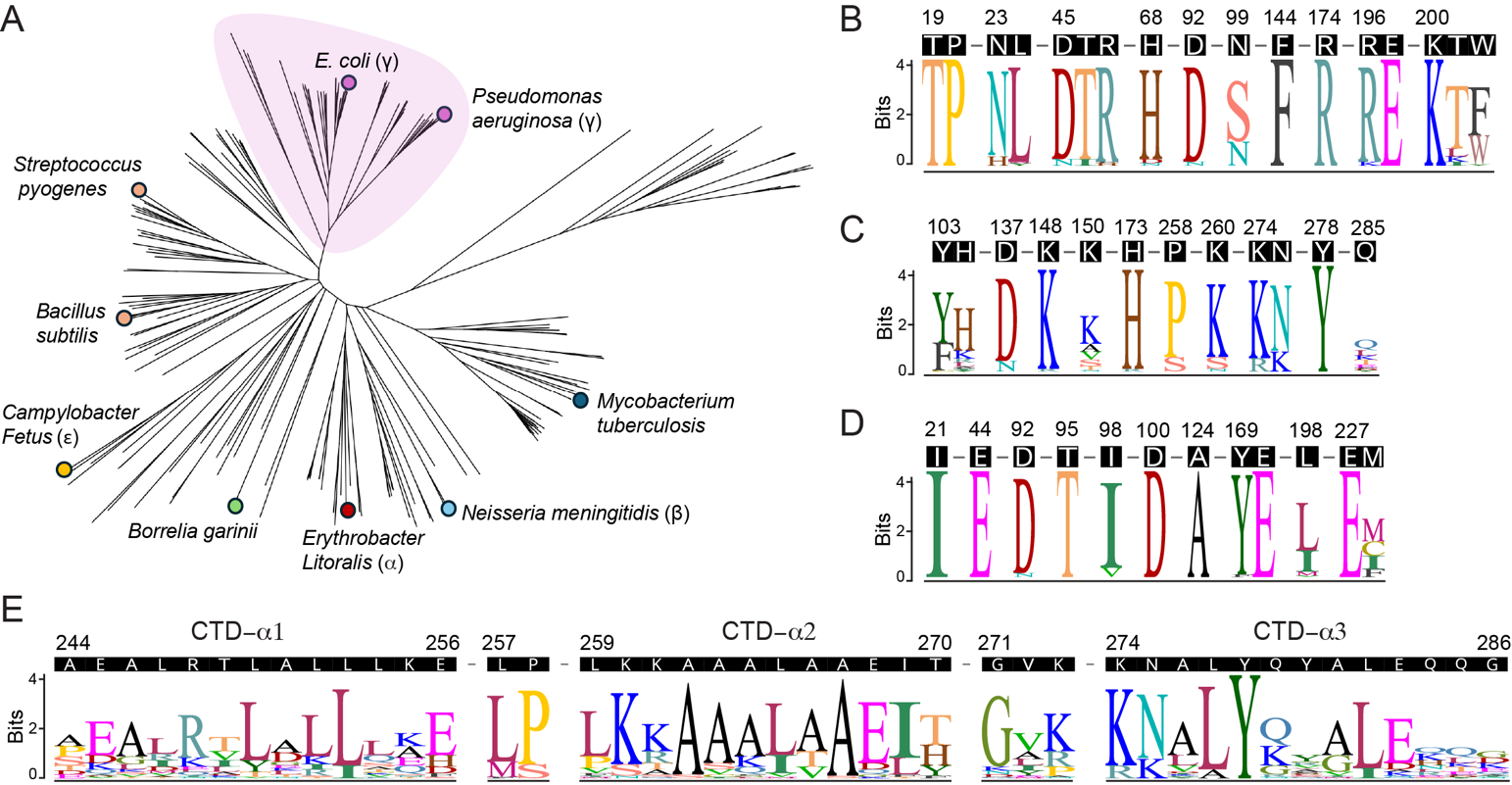

**Fig. S5. Conservation of RsmI among bacterial species and key functional residues among RsmI orthologs.** (*A*) Neighbor-joining phylogenetic tree displaying the distribution of RsmI among key bacterial families, with the Gammaproteobacterial RsmI clade highlighted (purple shading). Sequence logo of Gammaproteobacterial RsmI sequences showing propensities for (*B*) residues involved h44 interaction, (*C*) 30S platform contact, and (*D*) SAM binding and Mg^2+^ ion coordination. **E.** Sequence logo of Gammaproteobacterial RsmI sequences showing the residue conservation for the CTD domain. All sequence logo plots are accompanied by the *E. coli* RsmI sequence above them as reference.

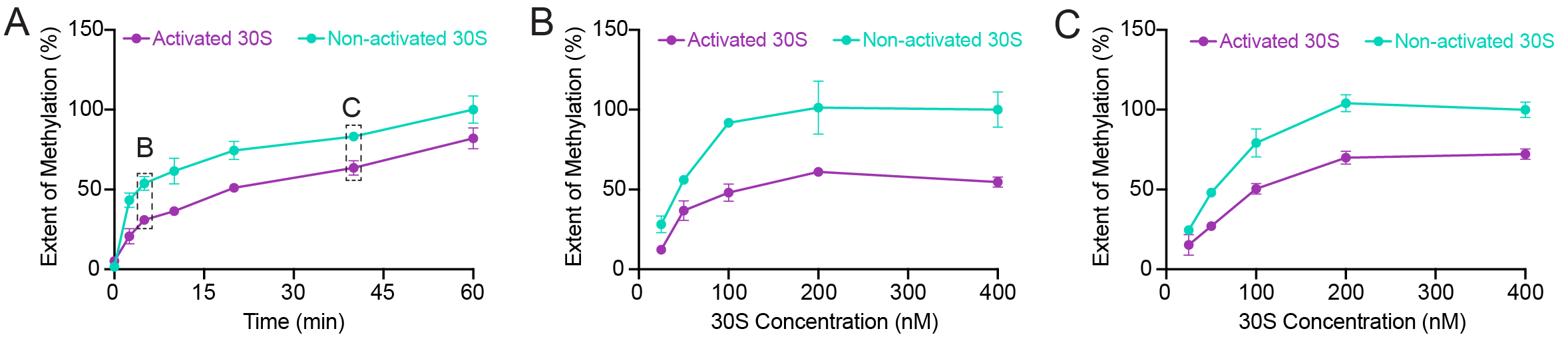

**Fig. S6. 30S subunit activation influences RsmI methylation activity *in vitro*.** (*A*) Time-course methylation assay for RsmI and Δ*rsmI* 30S under activating and non-activating conditions at a 1:1 enzyme to 30S ratio. Using the time-course as a guide, the impact of activation was compared by altering 30S subunit concentration at two time points: (*B*) early in the time course (5 minutes) and (*C*) near the reaction plateau (40 minutes) while keeping the enzyme concentration constant.

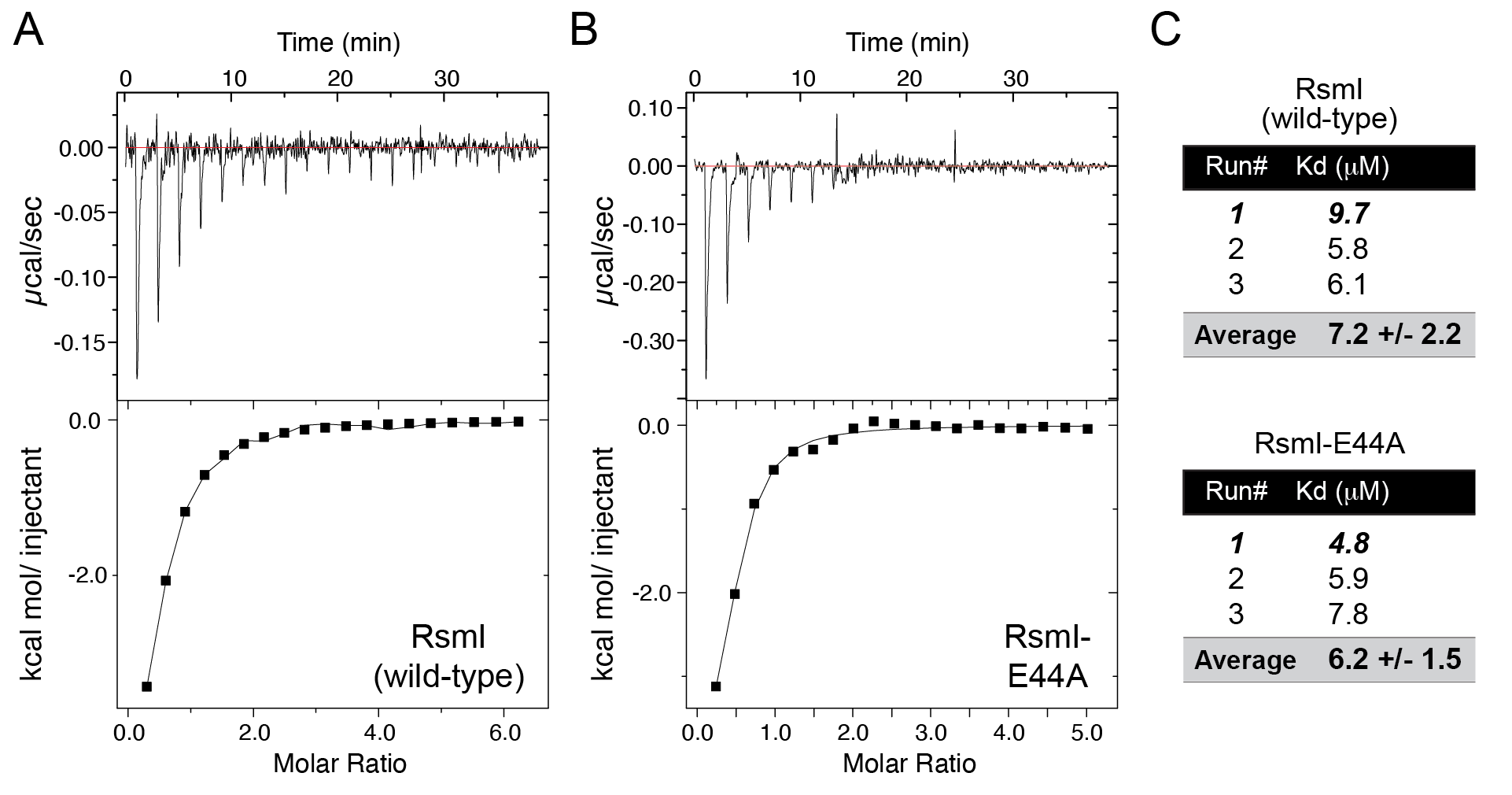

**Fig. S7. The E44A substitution does not impact SAM binding.** Representative ITC experiments (*top*, raw titration thermogram and *bottom*, integrated heats) for titration of SAM into (*A*) wild-type RsmI and (*B*) the RsmI E44A variant. (*C*) Individual and average K_d_ values obtained from the triplicate ITC experiments with each protein.

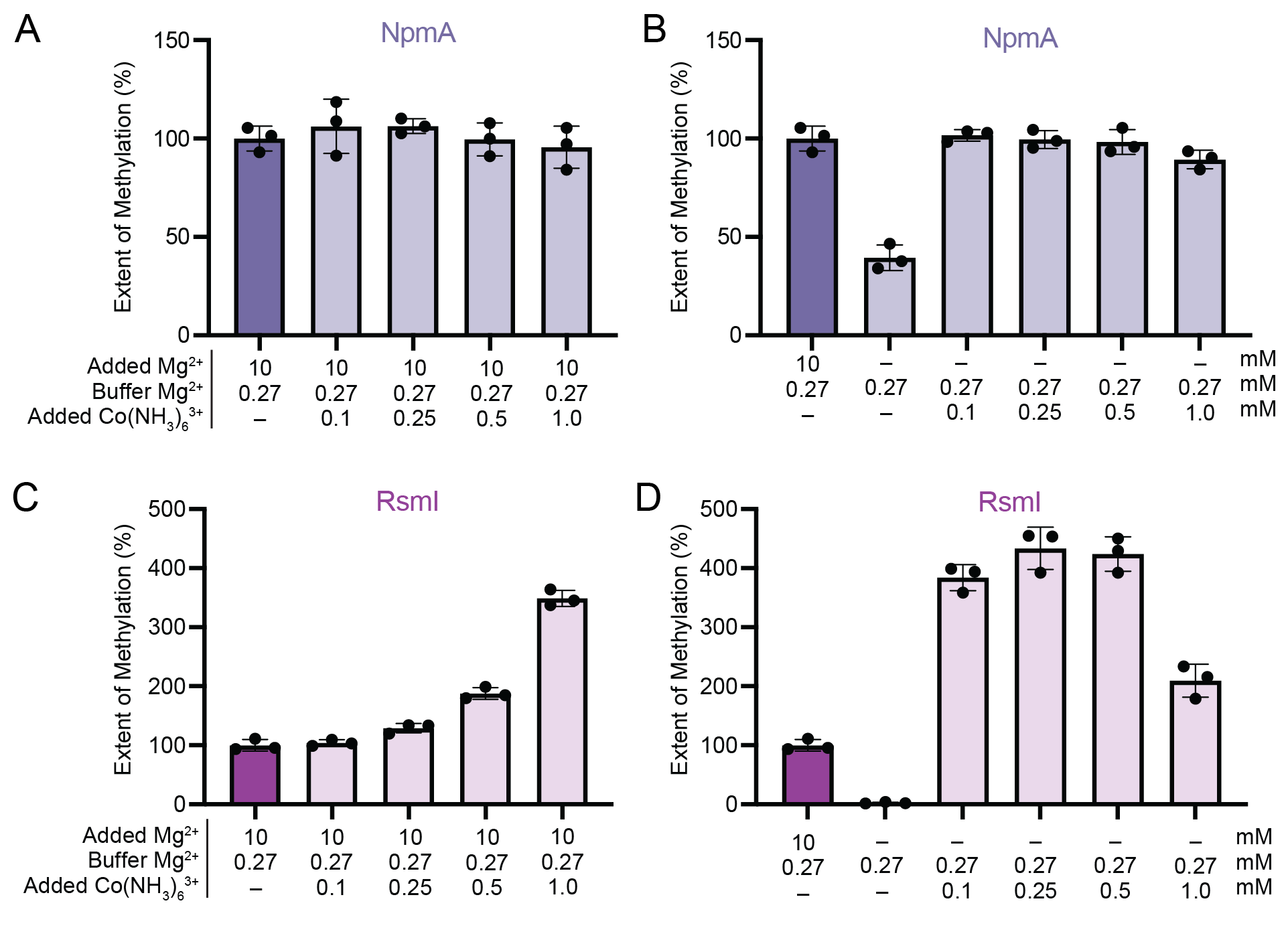

**Fig. S8: Methyltransferase assays with Co(NH_3_)_6_^3+^** **suggest that the divalent ion in the RsmI active site plays a predominantly structural role in coordinating C1402 for 2’-O-modification.** *In vitro* methyltransferase activity assays with Co(NH_3_)_6_^3+^ titrated from 0 to 1 mM for NpmA with (*A*) 10 mM Mg^2+^ or (*B*) no additional Mg^2+^ in the reaction buffer. (*C,D*) As for *panels A* and *B* but for RsmI.

**Supplementary Tables**

| **Table S1. Cryo-EM data collection, refinement, and model validation statistics** | | | |
| --- | --- | --- | --- |
|  | RsmI-30S subunit  (full complex) | RsmI-16S rRNA  (binding interface)**^a,b^** | 30S-Δ*rsmI*  (no RsmI)**^b^** |
| ***Deposition*** | | | |
| Coordinates (PDB) | 9PZG |  | 10FZ |
| Map (EMDB) | EMD-72071 |  | EMD-75144 |
| ***Data Collection*** |  |  |  |
| Magnification | ×105,000 |  |  |
| Voltage (kV) | 300 |  |  |
| Electron exposure (e^-^/Å^2^) | 49.27 |  |  |
| Defocus range (µm) | -0.4 to -2.9 |  |  |
| Pixel size (Å) | 0.41 |  |  |
| Symmetry imposed | C1 |  |  |
| Initial particle count | 3,781,314 |  |  |
| Final particle count | 74,553 | 264,656 | 58,273 |
| Map resolution (Å)  (FSC_0.143_, masked) | 2.42 | 2.55 | 2.91 |
| ***Refinement*** | | | |
| Model resolution (Å)  (FSC_0.143/0.5_, masked) | 2.43/2.92 | 2.57/2.92 | 2.91/3.21 |
| d_model | 2.70 | 2.90 | 3.00 |
| CC_mask_ | 0.84 | 0.87 | 0.79 |
| Model composition |  |  |  |
| Nonhydrogen atoms | 55,584 | 6,345 | 51,092 |
| Protein residues | 2,868 | 549 | 2,320 |
| Nucleotides | 1,529 | 95 | 1,530 |
| Ligands |  |  |  |
| NM6 (AN6) | 2 | 2 | – |
| Mg^2+^ | 79 | 1 | 77 |
| Water | 170 | 18 | 32 |
| ADP (B-factors) |  |  |  |
| Protein (min/max/mean) | 23.8/ 636.0/ 179.8 | 24.6/636.0/141.7 | 46.8/664.2/243.3 |
| Nucleotide (min/max/mean) | 20.0/430.4/159.4 | 20.0/220.1/92.8 | 40.7/980.9/244.6 |
| Ligand (min/max/mean) | 30.0/135.7/67.8 | 68.2/68.2/68.2 | 47.6/121.9/72.5 |
| Water (min/max/mean) | 10.0/515.0/85.9 | 10.0/515.0/119.6 | 66.8/1132.3/280.8 |
| R.M.S. deviations |  |  |  |
| Bond lengths (Å) | 0.004 | 0.010 | 0.002 |
| Bond angles (˚) | 0.764 | 1.287 | 0.459 |
| ***Validation*** | | | |
| MolProbity score | 1.89 | 1.48 | 1.89 |
| Clashscore | 6.84 | 6.82 | 7.89 |
| Rotamer outliers (%) | 1.66 | 1.38 | 0.05 |
| Ramachandran plot |  |  |  |
| Favored (%) | 95.0 | 98.3 | 92.6 |
| Allowed (%) | 5.0 | 1.7 | 7.3 |
| Outliers (%) | 0.0 | 0.0 | 0.0 |
| **^a^**Processing and refinement statistics for the focused map of the RsmI-30S subunit interface and corresponding truncated model comprising RsmI and its 16S rRNA tertiary surface (see Materials and Methods for further details).  ^b^Data collection parameters are identical to those for the RsmI-30S subunit (full complex) as the three structures were determined using the same image data set. | | | |

**Table S2. Conservation of select residues among RsmI family methyltransferases**

| Residue | Interaction | Residue identity (% occurrence)***^a^***^,^***^b^*** | |
| --- | --- | --- | --- |
|  |  | All | γ-Proteobacteria |
| T19 | A1408 | T (91.9) | T (100) |
| P20 | A1408 | P (92.4) | P (100) |
| I21 | SAM | I (82.3), L(11.6) | I (100) |
| N23 | C1407 | N (79.8), D (8.6), E (5.6) | N (90.7), H (9.3) |
| L24 | A1408 | L (69.2), V (6.1), I (4.0) | L (95.3), Y (2.3), M (2.3) |
| E44 | Mg^2+^ | E (100) | E (100) |
| D45 | C1400 | D (90.4), N (7.1) | D (93), N (7) |
| T46 | C1400 | T (87.9), A (3.5), E (3) | T (93), I (7) |
| R47 | C1400 | R (91.4), K (7.1) | R (95.3), H (2.3), F (2.3) |
| H68 | G1401 | H (71.7), G (7.6), F (6.1), | H (93) |
| D92 | A1408 | D (90.9), E (8.1) | D (95.3), N (4.7) |
| T95 | SAM | T (48.0), M (35.4), L (7.1) | T (100) |
| I98 | SAM | I (67.7), V (30.3) | I (90.7), V (9.3) |
| N99 | C1402 | S (81.3), A (8.6), N (8.1) | S (72.1), N (27.9) |
| D100 | SAM/ Mg^2+^ | D (100) | D (100) |
| Y103 | A790 | Y (41.4), F (17.7), A (15.7) | Y (62.8), F (34.9) |
| H104 | A790 | R (21.7), H (15.2), E (12.1), | H (67.4), K (11.6), R (7) |
| A124 | SAM | A (89.4), S (10.6) | A (100) |
| D137 | A790 | D (45.4), N (21.9), E (7.7) | D (88.4), N (11.6) |
| F144 | C1403 | F (91.9), Y (8.1) | F (100) |
| K148 | h24-h45 | K (78.3), D (6.1), E (4.5) | K (97.7), R (2.3) |
| K150 | h24-h45 | K (35), G (29.9), A (9.1) | K (51.2), A (14), V (14) |
| Y169 | SAM | Y (63.6), F (26.8), I (6.6) | Y (97.7), F (2.4) |
| E170 | SAM | E (96) | E (100) |
| H173 | h24-h45 | H (65.2), Y (17.7), R (5.6) | H (97.7), R (2.4) |
| R174 | C1403 | R (94.4), K (5.6) | R (100) |
| R196 | G1405 | R (86.9), K (5.6), A (4) | R (95.3), K (4.7) |
| E197 | C1404 | E (90.9), D (4.5), N (3.5) | E (100) |
| L198 | SAM | L (75.8), I (18.7) | L (100) |
| K200 | C1404 | K (100), L (4.5) | K (100) |
| T201 | C1407 | T (32.8), L (25.3), K (11.1) | T (79.1), L (9.3), K (7.0) |
| W202 | G1405 | F (33.3), H (25.3), Y (25.3), W (8.1) | F (60.5), W (37.2) |
| E227 | SAM | E (90.9), P (8.1) | E (100) |
| M228 | SAM | I (34.3), F (26.8), C (14.6) | M (39.5), I (23.3), C (23.3) |
| R248 | CTD (dimer) | R (30.7), A (11.7), L (11) | R (67.4), K (14) |
| L252 | CTD (dimer) | L (35.4), R (14.9), E (7.5) | L (65.1), K (11.6), R (9.3) |
| L253 | CTD (dimer) | L(41.5), I (21.4), V (15.1) | L(86), I (14) |
| L257 | CTD (dimer) | L (40.3), M (16.4), E (11.9) | L (79.1), M (18.6) |
| P258 | h23 | P (37.1), R (20.1), S (19.5) | P (83.7), S (16.3) |
| K260 | h23 | K (69.4), S (11.5), N (11.5) | K (83.7), S (14) |
| L265 | CTD (dimer) | L (28.8), E (14.1), Q (12.3) | L (69.8), I (23.3) |
| E268 | CTD (dimer) | K (32.7), E (24.7), A (16.7) | E (79.1), D (9.3) |
| I269 | CTD (dimer) | I (26.7), L (13.7), E (12.4) | I (72.1), L (23.3) |
| K274 | h23 | K (60.5), R (29.3) | K (86), R (14) |
| N275 | h23 | R (32.5), N (28.8), K (18.8) | N (76.7), K (23.3) |
| Y278 | h23 | Y (91.8), F (8.2) | Y (100) |
| Q285 | h23 | K (37.6), R (14.4), Q (13.6) | Q (33.3), L (16.7), K (13.9), T (13.9) |
| ***^a^***Residue percentages were calculated for the top three amino acids at each position, unless the third and fourth residues had identical frequencies, in which case all were included  ***^b^***Analysis was performed using all 198 RsmI sequences and a subset of 43 sequences from Gammaproteobacteria. | | | |

**Table S3. List of primers used in this study**

| Primer | Sequence 5’ to 3’ | Purpose |
| --- | --- | --- |
| *cat*-RsmI-Forward | **ATCGGCAATCTGGCGGATATCACCCAGCGTGCGTTAGAGG**CTGGTGTCCCTGTTGATACC | **Figure S1A, Step 1**: Generate *cat* cassette: 40 bp of *rsmI* sequence (bold/ underline) and pZS21 before and after *cat*. |
| *cat*-RsmI-Reverse | **TGCGGCCAGCGCCGCCGCTTTTTTCAGCGGCAGTTCTGCC**GCGTTTAAGGGCACCAATAAC |  |
| RsmI-ORF-Forward | GAAGATCTCAT**ATG**AAACAACACCAATCGGC | **Figure S1A, Step 3**: clone *rsmI* into pET-44 and verify *rsmI* deletion. Start/ end of *rsmI* ORF (bold/underline). |
| RsmI-ORF-Reverse | CTAAAGCTTTTA**TTA**CCCCTGCTGCTCC |  |
| pET-44-RsmI-Forward | GCAACCGCGAATTCACCTGTGG | ***rsmI* mutagenesis**:  Reverse primer for first-round PCR in MEGAWHOP contains the altered codon (bold/ underlined) |
| T19Y | CAGATTGCCGATTGG**ATA**CGGTACAATG |  |
| N23Y | GATATCCGCCAG**ATA**GCCGATTGG |  |
| E44A | CGAGTATC**TGC**GGCGGCAATC |  |
| T46A/R47A | GGTGTG**AGCAGC**ATCCTCGG |  |
| R47A | CCGGTGTG**AGC**AGTATCCTC |  |
| H68A | GTTGTTCGTT**AGC**GTCGTGCAG |  |
| D92A | GTTCCGGC**AGC**GGAAACCAG |  |
| Y103A | CACCAGATG**AGC**GCCAGGATC |  |
| Y103A/H104A | GTACGCACCAG**AGCAGC**GCCAGGATCGTTAATTAG |  |
| D137R | GTAACAGAAACG**CCG**AGAGGGTAAAC |  |
| F144A | GCAGGTAA**AGC**GCCTTCGTAAC |  |
| K148E | GGCCTTTTGA**TTC**GGCAGGTAAAAAG |  |
| K148E/K150E | GCC**TTC**TGA**TTC**GGCAGGTAAAAAG |  |
| K150E | GGCGGCC**TTC**TGATTTGG |  |
| H173A | GCTATCTAACAGACG**AGC**GGTAGATTC |  |
| R174A | GCTATCTAACAG**AGC**GTGGGTAGATTC |  |
| R174E | GCTATCTAACAG**CTC**GTGGGTAGATTC |  |
| R196A | GTCAGCTC**AGC**CGCCAGAAC |  |
| R196E | GTCAGCTC**TTC**CGCCAGAAC |  |
| K200A | GTTTCCCAGGT**AGC**GGTCAGCTC |  |
| K200E | GTTTCCAGGTTTCGGTCAGCTC |  |
| W202A | GTGAATGGTTTC**AGC**GGTTTTGGTC |  |
| E227A | CAATCAGCACCAT**AGC**GCCTTTGC |  |
| R248E | GCCAGCGT**TTC**CAGGGCATC |  |
| L253E | GTTCTGCCTG**TTC**CAGCGCCAG |  |
| P258G | GCTTTTTTCAG**ACC**CAGTTCTGC |  |
| K260E | CCGCTTT**TTC**CAGCGGC |  |
| Y278A | CGCATACTT**AGC**CAGCGCATTTTTC |  |
| ΔCTD-α_3_ | GCATTTTTCTTCAC**TCA**GTGAATTTCTGCG |  |
| ΔCTD | GTCTTCTTCCTG**TCA**TTTATGACCTTCG |  |
